# Transcription terminator-mediated enhancement in transgene expression in maize: preponderance of the AUGAAU motif overlapping with poly(A) signals

**DOI:** 10.1101/2020.06.08.140475

**Authors:** Po-Hao Wang, Sandeep Kumar, Jia Zeng, Robert McEwan, Terry R. Wright, Manju Gupta

## Abstract

The selection of transcription terminators (TTs) for pairing with high expressing constitutive promoters in chimeric constructs is crucial to deliver optimal transgene expression in plants. In this study, the use of the native combinations of four polyubiquitin gene promoters and corresponding TTs resulted in up to >3-fold increase in transgene expression in maize. Of the eight polyubiquitin promoter and TT regulatory elements utilized, seven were novel and identified from the polyubiquitin genes of *Brachypodium distachyon, Setaria italica*, and *Zea mays*. Furthermore, gene expression driven by the Cassava mosaic virus promoter was studied by pairing the promoter with distinct TTs derived from the high expressing genes of *Arabidopsi*s. Of the three TTs studied, the polyubiquitin10 gene TT produced the highest transgene expression in maize. Polyadenylation patterns and mRNA abundance from eight distinct TTs were analyzed using 3’-RACE and next-generation sequencing. The results exhibited one to three unique polyadenylation sites in the TTs. The poly(A) site patterns for the StPinII TT were consistent when the same TT was deployed in chimeric constructs irrespective of the reporter gene and promoter used. Distal to the poly(A) sites, putative polyadenylation signals were identified in the near-upstream regions of the TTs based on previously reported mutagenesis and bioinformatics studies in rice and *Arabidopsis*. The putative polyadenylation signals were 9 to 11 nucleotides in length. Six of the eight TTs contained the putative polyadenylation signals that were overlaps of either canonical AAUAAA or AAUAAA-like polyadenylation signals and AUGAAU, a top-ranking-hexamer of rice and *Arabidopsis* gene near-upstream regions. Three of the polyubiquitin gene TTs contained the identical 9-nucleotide overlap, AUGAAUAAG, underscoring the functional significance of such overlaps in mRNA 3’ end processing. In addition to identifying new combinations of regulatory elements for high constitutive trait gene expression in maize, this study demonstrated the importance of TTs for optimizing gene expression in plants. Learning from this study could be applied to other dicotyledonous and monocotyledonous plant species for transgene expression.

## 1. Introduction

The use of varying transcription terminators (TTs), containing the 3’ untranslated regions (3’ UTRs), in chimeric constructs has resulted in multifold increases in gene expression in transgenic plants. TT-mediated increases in gene expression were reported in transgenic tobacco (Ingelbrecht et al., 1989; Richter et al., 2000; Diamos and Mason, 2018), potato (An et al., 1989), tomato (Hiwasa-Tanase et al., 2011), *Arabidopsis* and rice (Nagaya et al., 2009). These studies illustrated the versatility of TTs for optimizing gene expression in transgenic plants.

The mRNA biogenesis entails 5’ end capping, intron splicing, and 3’ end formation of pre-mRNA prior to its export from the nucleus into the cytoplasm. All three steps occur coordinately with high specificity and efficiency [reviewed in (Wahle and Keller, 1992; Proudfoot et al., 2002; Proudfoot, 2004)]. The 3’ end of the mRNA cross-talks with the 5’ end through RNA “looping” and stimulates transcription initiation through coordinated recycling of RNA polymerase II. This process is integral to efficient and sustained transcription of mRNA (Mapendano et al., 2010; Andersen et al., 2012). Similarly, already assembled polyadenylation protein complexes at the 3’ end enhance 3’ end formation (O’Sullivan et al., 2004; Tan-Wong et al., 2009; Andersen et al., 2012; Tan-Wong et al., 2012).

The TTs present downstream of the gene-of-interest contain polyadenylation [poly(A)] signals, which control the steps involved in 3’ end formation: recognition, endonucleolytic cleavage, and polyadenylation of primary RNA (pre-mRNA). These steps impact gene expression by influencing mRNA termination, stability, localization, export to cytoplasm and translation efficiency. Dissection of the poly(A) signals with conventional mutagenesis studies [reviewed in (Hunt, 1994)] identified three main *cis* elements in 3’ UTR requisite for 3’ end processing, which were further confirmed through the genomic scale sequencing of transcripts in *Arabidopsis* (Loke et al., 2005) and rice (Shen et al., 2008). These are far-upstream elements (FUEs) and near-upstream elements (NUEs) located upstream of the cleavage site (CS). The FUE and NUE are U- and A-rich, respectively. The CS of ∼7 nucleotides (nt) in length is embedded in U-rich elements (URE), with ∼10 nt on each side of the CS. The combination of the CS and URE is referred to as the cleavage element (CE) (Loke et al., 2005). The CS has a characteristic YA sequence (Y = U or C) immediately upstream of the pre-mRNA cleavage. The FUE impacts the function of the NUE, which in turn controls the function of each of the CSs and poly(A) sites in a transcription unit (Hunt, 1994; Rothnie et al., 1994). The NUE extends up to 30-35 nt upstream of the CS while the FUE spans up to 135-150 nt depending on the plant species. Highly conserved in the NUE of mammalian genomes, a canonical AAUAAA polyadenylation motif is consistently observed within 10-30 nt from the CS in *Arabidopsis* (Loke et al., 2005) and 10-35 nt in rice (Shen et al., 2008). However, this sequence is present in only 10% of the *Arabidopsis* and 7% of rice transcripts (Loke et al., 2005; Shen et al., 2008). Studies show AAUAAA-like polyadenylation motifs in plants are generally tolerant to 1- or 2-nt mutations without affecting the polyadenylation efficiency in plants (Rothnie et al., 1994; Li and Hunt, 1995). Only complete deletion of this sequence has been shown to abolish the polyadenylation (Wu et al., 1995). The *cis*-acting elements of the poly(A) signal and *trans*-acting polyadenylation complexes that recognize and bind to these *cis* elements are involved in 3’ end formation of the pre-mRNA. Polyadenylation complex studies in *Arabidopsis* led to the identification of up to 28 relevant protein factors (Hunt, 2008).

There are single to multiple poly(A) sites corresponding to 3’ UTRs in plants. These sites contribute to alternative polyadenylation (APA). Accordingly, one gene can give rise to multiple distinct mRNA isoforms with varying degrees of stability, coding potential, and gene regulation capability (Ji and Tian, 2009; Lutz and Moreira, 2011; Hunt, 2012). Results from genome-wide studies showed that TTs of plant genes are especially rich in APA sites, and up to 60% of *Arabidopsis* genes and 47%-82% (based on the technique deployed for mRNA sequencing) of rice genes utilize APA (Meyers et al., 2004; Di Giammartino et al., 2011; Shen et al., 2011; Wu et al., 2011; Hunt, 2012). The length of mRNA is affected depending on the APA site recognition in 3’ UTR. The longer mRNA may encompass micro RNA (miRNA) binding sites, RNA binding protein factors, and AU-rich elements that can regulate gene expression (Barreau et al., 2005; Fabian et al., 2010).

In plants and animals, APA is affected by physiological conditions (such as cell growth, differentiation, and development), pathological events (such as cancer), and environmental cues (Ji and Tian, 2009; Mayr and Bartel, 2009; Di Giammartino et al., 2011). APA could be due to tissue-specific expression of the polyadenylation-related genes, which suggest different polyadenylation complex organization in those tissues (Hunt, 2008). Other well-studied examples in plants include APA in the TTs of genes of autonomous flowering time pathway, including *FCA, FPA and FLOWERING LOCUS C (FLC)* in *Arabidopsis* (Simpson et al., 2003; Hornyik et al., 2010; Liu et al., 2010).

Gene silencing is frequently observed in transgenic plants when high gene expression is driven by strong constitutive or viral promoters (Bass, 2000; Baulcombe, 2004; Luo and Chen, 2007). In this particular mechanism, aberrant un-polyadenylated mRNA is synthesized, and it is distinguished from the canonical polyadenylated mRNA through an unknown mechanism (Baeg et al., 2017). The aberrant mRNA is then converted into double-stranded RNA through the activity of RNA-dependent-RNA-polymerase6 (RDR6). Subsequently, the double-stranded RNA gets converted into small interfering RNA (siRNA), a molecule fundamental to gene silencing (Bass, 2000; 2002). A limited number of 3’ UTRs of *Agrobacterium* or viral origins, including *nopaline synthase* (*nos*), *octopine synthase* (*ocs*) and cauliflower mosaic virus 35S (CaMV 35S), have been commonly utilized as regulatory elements (REs) in gene expression cassettes of transgenic plants (Diamos and Mason, 2018). However, these REs did not yield expression in the expected range (Rose and Last, 1997; Luo and Chen, 2007; Diamos and Mason, 2018). When paired with strong constitutive or viral promoters, these 3’ UTRS have been shown to produce improperly terminated and un-polyadenylated transcripts. This results in production of transcripts with unprotected 3’ ends, which are substrates for RDR6 and trigger gene silencing (Baulcombe, 2004; Luo and Chen, 2007). Gene silencing in the transgenic products could make it less efficacious in the field.

First-generation biotech crops mainly involved simple traits. Next-generation products attempt to deliver crop improvements by expressing multiple traits consisting of insect resistance, herbicide tolerance, and other transgenic traits (Que et al., 2010). Since each transgene usually requires a unique promoter and TT pair for expression, multiple REs are required to express different transgenes within one molecular stack. This frequently leads to repeated use of the same promoter and TT combination due to limited availability of verified REs. Multi-gene constructs driven by the same promoter could also lead to gene silencing (Peremarti et al., 2010). While tissue-specific promoters are desirable for certain traits, strong and constitutive expressing promoters are required for the majority of insect and herbicide traits. Promoters from viral (Verdaguer et al., 1998; Schenk et al., 1999; Bohorova et al., 2001; Samac et al., 2004; Davies et al., 2014) and plant polyubiquitin (UBQ) genes [summarized in (Lu et al., 2008; Mann et al., 2011)] have been used to obtain enhanced constitutive transgene expression.

Ubiquitin is a small conserved protein of 76 amino acids found in all eukaryotes (Sullivan et al., 2003). The promoters, introns, 5’ UTRs and 3’ UTRs of these UBQ genes are quite diverse, in comparison to the highly conserved protein sequences. Given the high constitutive expression of UBQ genes in the majority of plant tissues, UBQ promoters and TTs are very attractive for plant biotechnological applications. The release of whole-genome sequences from different plant species provided the opportunity to clone and study new UBQ promoters and respective TTs from other monocot species. In this report, we describe novel UBQ promoters and TTs derived from monocot species. Twelve combinations of promoters (UBQ and viral) and TTs (UBQ and *Arabidopsis*) were evaluated in maize for transgene expression. Analysis of gene silencing, identification of poly(A) sites and corresponding putative poly(A) signals were performed to evaluate consistency and sustainability of transgene expression in maize and better understanding of functional *cis* elements in the TTs.

## 2. Materials and Methods

### 2.1. Identification of promoters and transcription terminators

*Z. mays polyubiquitin1* (*ZmUbi1*) gene coding sequence, GRMZM2G409726 (B73 genotype; Christensen et al., 1992), of the maize genome database was BLASTx searched against the *Brachypodium distachyon* and *Setaria italica* genomes in the Phytozome database (http://www.phytozome.net) (Goodstein et al., 2011). Two matching sequences from *B. distachyon* (Bradi1g32460, referred to as *BdUbi-C*; and Bradi1g32700, referred to as *BdUbi1*), and one from *S. italica* (Seita.1G078900, referred to as *SiUbi2*) were identified. Putative promoters containing the transcription start site, upstream elements, exon and intron sequences (∼2 kb DNA sequence upstream from the predicted translational start site [ATG]), and putative TTs (∼1 kb DNA sequence downstream of the predicted translational stop site [TAA, TAG or TGA]) were identified (Kumar and Worden, 2017). Similarly, 910 bp of the putative TT sequence from the *ZmUbi1* gene was identified in the maize database, PCR-amplified, and cloned (Kumar et al., 2017b) to pair with the existing *ZmUbi1* promoter (Christensen et al., 1992). These sequences were used for cloning and transgene expression studies described below.

### 2.2. Cloning and construction of plant transformation vectors

Gateway (Thermo, NY, USA) entry and destination vectors were constructed by standard molecular cloning methods [see (Kumar et al., 2015; Kumar and Worden, 2017)]. Three promoter and three TT sequences derived from the *BdUbi1, BdUbi1-C* and *SiUbi2* genes (Supplementary Figure 2A-2C) were synthesized (GeneArt, InVitrogen, CA, USA). These sequences were flanked by 15-18 nucleotide homology fragments on both ends for seamless cloning (Invitrogen, CA, USA) based on DNA recombination technology (Zhu et al., 2007; Kumar et al., 2017a). Type II restriction enzyme sites were inserted outside of the 15-18 nucleotide homology for isolation of RE or gene fragments. Seamless cloning compatible fragments of the following DNA sequences were obtained by PCR: *ZMUbi1* promoter and TT; two reporter genes, yellow fluorescent protein gene [*phiyfp*, referred to as *yfp* (Evrogen, Moscow, Russia) (Shagin et al., 2004)] and *cry34Ab1*gene [derived from *Bacillus thuringiensis* (Moellenbeck et al., 2001)]; potato protease inhibitor II (*StPinII*) gene TT; Cassava mosaic virus (CsVMV) promoter (Verdaguer et al., 1998); and three TTs derived from *Arabidopsis thaliana* genes, *elongation factor 1* (*AtELF1* or *AtELF1*α; Gene ID: AT1G07920), *ubiquitinC9* (*AtUBC9;* Gene ID AT4G27960), and *polyubiquitin-10* (*AtUbi10;* Gene ID: AT4G05320). The *Arabidopsis* genes, *AtELF1, AtUbiC9* and *AtUbi10* have been shown to express at high levels in *A. thaliana* using qRT-PCR (Czechowski et al., 2005). The *yfp* gene was interrupted by a second intron of 189 bp of the ST-LS1 gene (Vancanneyt et al., 1990). The *cry34Ab1* [also a component of SmartStax™ (Siebert et al., 2012)] and *yfp* genes were selected as reporters because of the availability of high-quality quantitative ELISA for determination of protein expression.

Expression constructs for *Agrobacterium*-mediated maize transformation were assembled by using the Gateway recombination reactions that employed a destination binary vector pDAB109805 and entry vectors as described above. The destination vector pDAB109805 contained the *aryloxyalkanoate dioxygenase* (*aad-1*) herbicide tolerance gene (Wright et al., 2010) controlled by a sugarcane bacilliform virus (SCBV) promoter (Samac et al., 2004) and TT from the maize lipase gene (ZmLip1) (Cowen et al., 2007). The Gateway recombination reaction was used to recombine entry vectors with the destination vector to obtain expression vectors (Figure 1). The reporter cassette was located 5′ to the *aad-1* cassette, and separated by a spacer sequence that ranged in size from 278 bp to 312 bp depending on the promoter tested. Two positive control constructs, *ZmUbi1* promoter/*yfp*/*StPinII* TT (121) and *ZmUbi1*/*cry34Ab1*/*StPinII* TT (746), were assembled to use as benchmarks for gene expression evaluations. More details of *yfp* and *aad-1* are described previously (Beringer et al., 2017). The *aad-1* cassette was the same in all constructs. All binary constructs were transformed into a *recA*-deficient EHA105 *Agrobacterium* strain for maize transformation as described previously (Merlo et al., 2012). Bacterial colonies were isolated, and plasmid DNA was extracted and validated via restriction enzyme digestion and sequencing.

**Figure 1.**
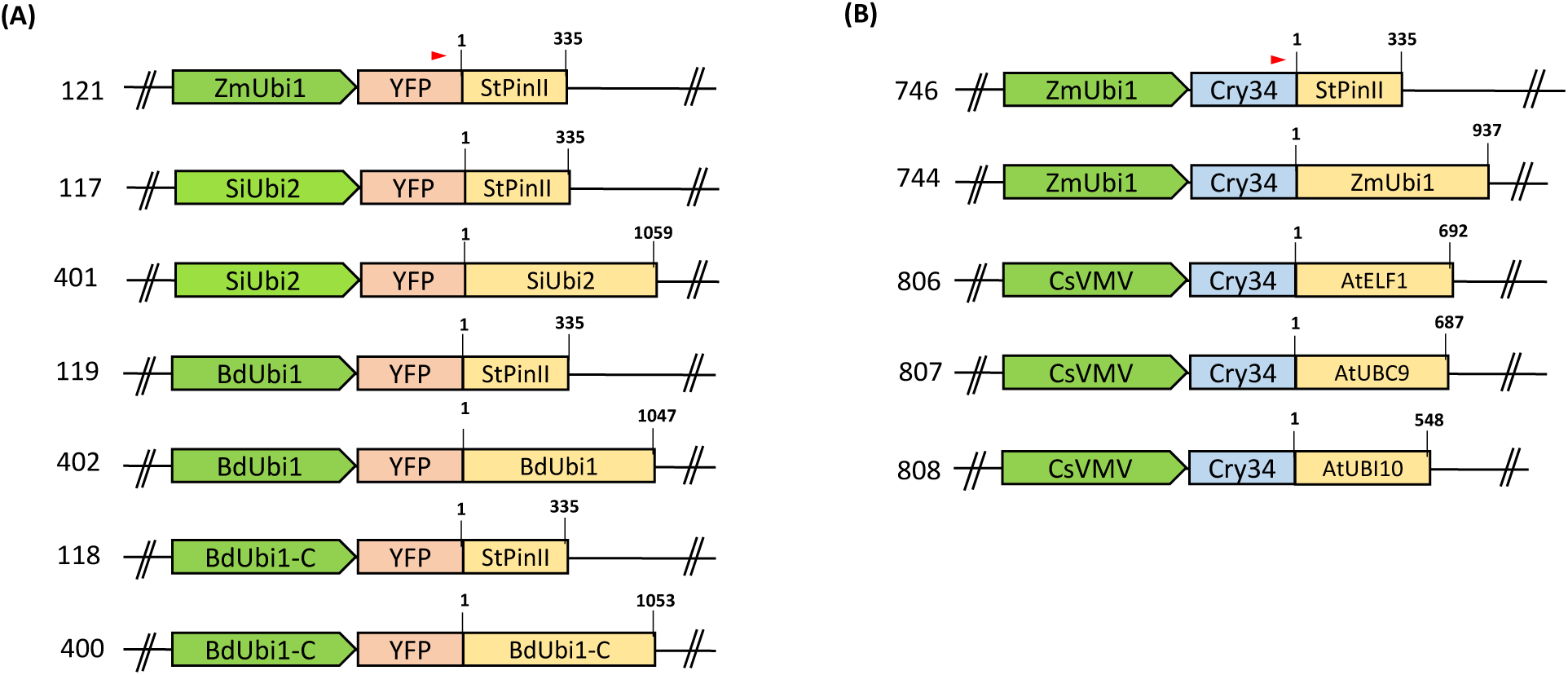
Schematic representation of assembled T-DNA cassettes: promoters and TTs are depicted as green arrows and yellow boxes, respectively. Distinct RE combinations were tested using the *yfp* (A) and *cry34Ab1* (B) reporter genes as depicted by pink and blue boxes, respectively. The T-DNA backbone and the selection marker, *aad-1*, present at the 3’ end of the cassettes are not shown in the schematic. The numbers on the left of the cassettes denote the construct numbers. The numbers on the top left of the TT boxes depict coordinates corresponding to the 3’ RACE sequencing data and that on the top right of the boxes denote the total length of the TTs. The red arrow on top of the reporter gene box displays the oligo position for 3’ RACE analysis.

**Figure 2.**
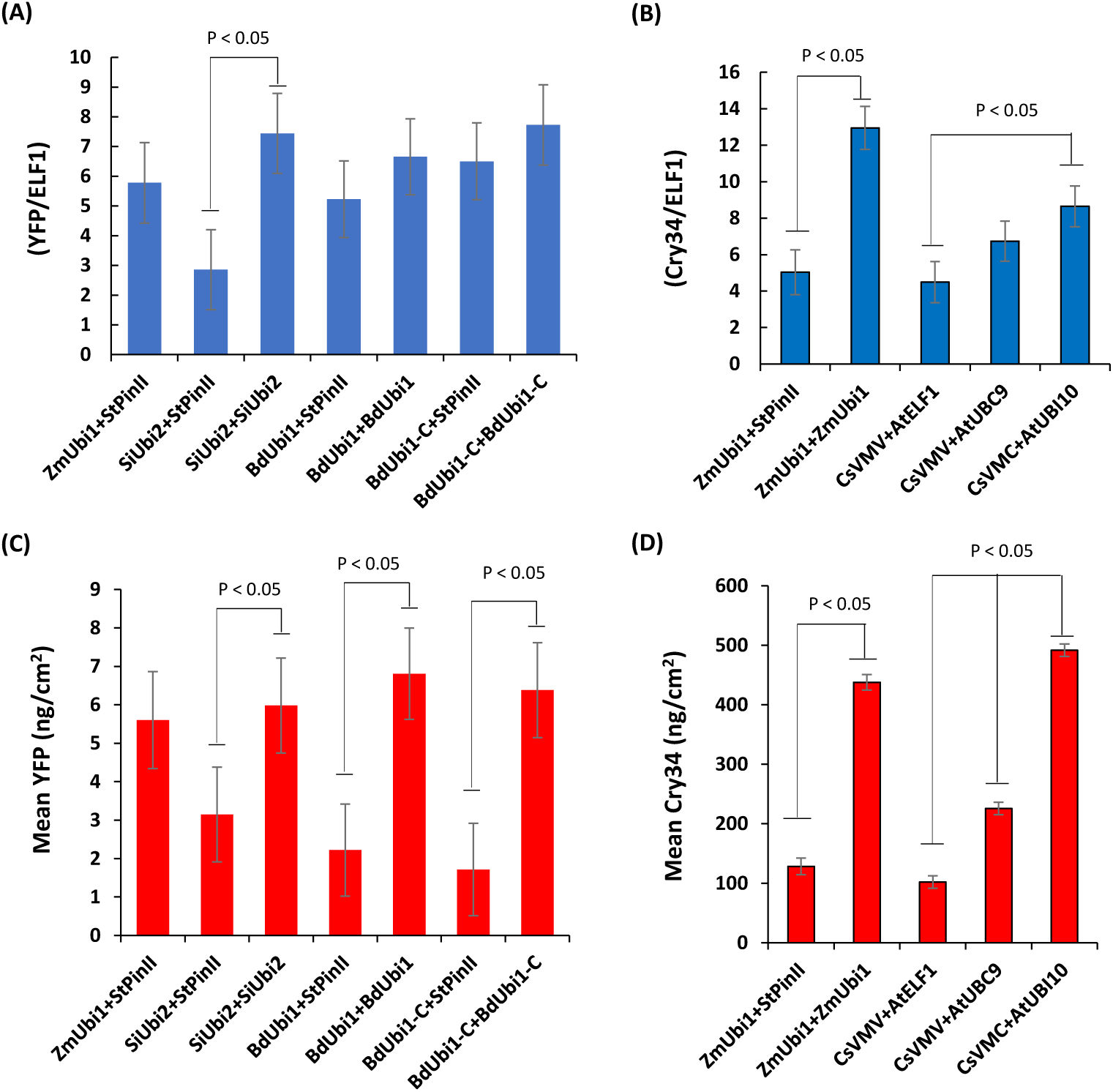
Transgene expression analysis of mRNA and protein in T_1_ plants containing the *yfp* and *cry34Ab1* reporter genes. qRT–PCR analysis of *yfp* (A) and *cry34Ab1* (B) for relative mRNA transcript levels in V6 leaves of transgenic maize plants. The same plants were used to quantitate protein accumulation for *yfp* (C) and *cry34Ab1* (D). All plants were derived from the hemizygous T_1_ plants of distinct transgenic events. For each construct, the data points were collected from at least three individual transgenic events and minimum 10 sibling plants per event at V6. Data are represented as mean ± SD of the combined data points for each construct, and differences were analyzed with Student’s t-test. The comparison showing a significant difference is indicated (p<0.05). The negative control construct (121 and 746) transgenic plants when used with Figure 1B and Figure 1A group of transgenic plants, respectively, did not show any mRNA or protein expression.

### 2.3. Maize transformation and plant growth conditions

*Agrobacterium*-mediated transformation of maize and plant growth conditions were described previously (Beringer et al., 2017). T_0_ plants were assayed for *yfp* or *cry34Ab1*, and *aad-1* copy number using quantitative PCR assays (see below). One-to two-copy events (plants with one-to two-copy insertion of the transgenes into the maize genome) were crossed to c.v. B104 non-transgenic transformation lines to obtain T_1_ seed. The T_1_ hemizygous seeds were grown in the green-house to the V2/V3 growth stage (Abendroth et al., 2009) and one leaf punch was sampled for reconfirmation of copy number for the reporter and *aad-1* genes. Subsequently, all plants were sprayed with a selective trea™ent of Assure II (quizalofop) herbicide spray to kill *aad-1* null plants (Wright et al., 2010). Events that segregated as 1:1 for *aad-1* positive *vs*. null plants and contained one-to two-copies of reporter gene were used in mRNA and protein gene expression analyses.

### 2.4. Quantitative PCR for copy number determination

Putative transgenic maize plants were sampled at the V2/V3 stage for transgene copy number estimation using quantitative PCR. Total DNA was extracted from the leaf samples, using MagAttract^®^ DNA extraction kit (Qiagen, MD, USA) as per the manufacturer’s instructions. Copy number assays for *yfp* and *aad-1* were performed as described previously (Kumar et al., 2015; Beringer et al., 2017). The *cry34Ab1* quantitative PCR assays were performed similarly except for the following details. The following primers and probes were used: *cry34Ab1* (Forward primer: GCCAACGACCAGATCAAGAC; Reverse primer: GCCGTTGATGGAGTAGTAGATGG and Probe: FAM-CACTCCCCACTGCCT-MGB). The invertase gene (Genebank Accession no. U16123.1) was used as an endogenous reference control, and the same primers and probes for this gene were used as described previously (Beringer et al., 2017). DNA was analyzed in a Roche LightCycler^®^ 480 System under the following conditions: 1 cycle of 95°C for 10 min; 40 cycles of the following 3-steps: 95°C for 10 seconds, 58°C for 35 seconds, and 72°C for 1 second; and a final cycle of 4°C for 10 seconds. *Cry34Ab1* copy number was determined by comparison of Target (gene of interest)/Reference (Invertase gene) values for unknown samples (output by the LightCycler^®^ 480) to Target/Reference values of *cry34Ab1* copy number controls.

### 2.5. Protein quantification with ELISA

Protein extractions from T_1_ maize leaves at the V6 stage were completed for all samples as described previously (Beringer et al., 2017). YFP and Cry34Ab1 ELISA assays were carried out as described previously (Kumar et al., 2015).

### 2.6. RNA extraction and RT__J ***qPCR analysis for transcript quantitation***

Quantitation of mRNA expression was completed as described previously (Wang et al., 2018). Briefly, plant RNA extractions were carried out using the Norgen Plant/Fungi RNA purification kit (Norgen Biotek, ON, Canada) using four leaf punches of maize leaves at V6. Total RNA was extracted from freeze-dried leaves and the first-strand cDNA synthesis was performed using a high capacity cDNA RT kit (ThermoFisher Scientific, USA) with random hexamers and Multiscribe™ reverse transcriptase following the manufacture’s protocol.

The resultant cDNA was quantitated via qRT-PCR with a hydrolysis probe. The assays were run for the target gene (i.e., *cry34Ab1* and *yfp*) and the *Z. mays elongation factor 1* (*elf1)* gene as the reference gene in biplex reactions. The assay was set up at 1× final concentration in a 10 µL volume containing 0.4 µM of each primer and 0.2 µM of each probe. Primers and probes for *yfp* and *cry34Ab1* were the same as described for copy number determination. Primers and probes for *elf1* were as follows: (Forward primer: ATAACGTGCCTTGGAGTATTTGG; Reverse primer: TGGAGTGAAGCAGATGATTTGC and Probe: Vic-TTGCATCCATCTTGTTGC-MGB). A two-step qRT-PCR program was performed for 40 cycles: denaturation at 95 °C for 10 s and extension at 58 °C for 40 s concomitant with fluorescence acquisition. Cp values were calculated based on the point at which the fluorescence signal crossed the background threshold using the fit points algorithm (LightCycler® software release 1.5). The relative transcript level was calculated using 2^ (−ΔCt), where ΔCt = Ct (target gene) - Ct (reference gene).

### 2.7. Determination of polyadenylation sites in transcription terminators

The poly(A) sites in TTss were determined using a 3’ RACE system (ThermoFisher Scientific, CA, USA) that amplified gene-specific 3’ ends from mRNA followed by sequencing of the resulting cDNA. Total RNA used for transcript quantitation (described above) was utilized for 3’ RACE analysis. Briefly, the polyA^+^ mRNA samples were converted into cDNA using SuperScript II reverse transcriptase and an oligo-dT adapter primer in accordance with the manufacturer’s instructions. After the conversion of the first-strand cDNA, the RNA template was removed from the cDNA-RNA hybrid molecules by digestion with RNase H to increase the sensitivity of PCR. Subsequently, gene-specific cDNA was amplified by PCR using a gene-specific primer that annealed to the 3’ end region of *cry34Ab1* (TCGACGTGAACAACAAGACC) or *yfp* (AAGAGCGCACAATCACCTTT) exon, and an adapter primer that targeted the poly(A) tail region of the first-strand cDNA under the following conditions: 1 cycle of 95°C for 5 min; 20 cycles of the following 3-steps: 95°C for 30 seconds, 62°C for 30 seconds, and 72°C for 90 second; and a final cycle of 72°C for 5 min. The PCR fragments were cleaned via a Gel DNA recovery kit (Zymo, CA, USA) and subjected to Next Generation Sequencing (NGS).

### 2.8. Protocol for NGS sequencing

One µg of 3’ RACE amplicons were diluted to 20 ng/µl in elution buffer and used as input in a modified KAPA Hyper Prep library production protocol (Kapa Biosystems, MA, USA) to generate libraries for Illumina sequencing. Since amplicon sizes ranged from about 100 – 800 bp, the fragmentation step in the protocol was skipped. End repair and addition of a single adenine (A) extension was completed as instructed, followed by ligation to indexed adapters overnight at 4°C. A 0.8X AxyPrep (Thermo Scientific, MA, USA) post-ligation bead cleanup was carried out with final elution in 20 µl of TE buffer. Ligated products were enriched with PCR under the following conditions: initial denaturation at 98°C for 45 s; 4 cycles of 98°C for 15 s, 60°C for 30 s, and 72°C for 30 s; and a final extension at 72°C for 1 min. PCR products were subsequently cleaned with AxyPrep beads at 1X concentration and eluted in 30 µl of TE buffer. Final library products were assessed for quality on LabChip GX using the DNA High Sensitivity Reagent Kit (Perkin Elmer, MA, USA), diluted to 2 nM concentration and pooled. This was followed by denaturation with sodium hydroxide and further dilution to 6 pM in hybridization buffer for loading into a MiSeq system. Sequencing was set up in accordance with Illumina’s recommended protocol for an indexed 100-cycle paired-end run.

### 2.9. 3’ RACE data analysis

NGS reads obtained from 3’ RACE amplicons were first trimmed against the adapter sequence to get high quality reads. They were then mapped to corresponding reference construct sequences by using Bowtie (version 2.3.2) (Langmead et al., 2009) with the following parameter settings: “--local --no-discordant --no-unal --no-mixed -X 5000”. After the comprehensive mapping, a custom script was used to generate the distribution of the mapped reads. The locations of the 3’ end of the mapped reads were extracted and summed up to obtain the raw counts for transcription termination sites. The raw counts were further normalized by the sequencing depth to allow comparison between different sites and different samples.

### 2.10. Small RNA library construction, sequencing, and bioinformatics analyses

Leaf samples collected for mRNA quantitation, as described above, were used in small RNA (sRNA) analysis for gene silencing. Total RNA was isolated using TRI Reagent Solution as per the manufacturer’s instructions (Ambion, Inc., USA), with an additional step of extraction with acid phenol (pH 4.5). RNA was quantified using a NanoDrop (Thermo Scientific, MA, USA). The quality of total RNA was evaluated using LabChipGX (Caliper LifeSciences, Waltham, MA, USA) and total RNA was used for sRNA library preparation and sequencing. The sRNA library was prepared, and data analyzed as described previously (Sidorenko et al., 2017). Raw sequencing reads were first trimmed of adapter sequences and then mapped to the transgene sequences. Briefly, Bowtie 2 (version 2.3.2) (Langmead et al., 2009) was used to map the trimmed and cleaned sRNA reads to a combined genome reference, including maize genome and transformed construct, with the following criteria: 1) no mismatches, and 2) only randomly reporting one location if a read could be mapped to multiple positions (all other parameters taking default values). The number of reads mapped to a specific gene or region was calculated by Samtools (version 0.1.9) (Li et al., 2009) then normalized by RPM (reads number per million). Another in-house script was used to do strand selection and to calculate the distribution of sRNA length. Finally, IGV (Integrative Genomics Viewer: version 2.5.3) was used to visualize the mapping results. sRNA abundance was calculated per one million genome□matched reads (RPM).

## 3. Results

### 3.1. Identification of promoters and transcription terminators

Alignment of the *ZmUbi1* gene coding sequence with that of the *SiUbi2, BdUbi1*, and *BdUbi1-C* revealed 100% amino acid homology in the basic 76 amino acid repeat unit (Supplementary Figure 1). However, the *BdUbi1* and *BdUbi1-C* polyubiquitin gene coding sequences contained pentameric repeats of the 76 amino acids each instead of the heptameric repeats characteristic of the *ZmUbi1*. The *SiUbi2* polyubiquitin gene coding sequence also contained the heptameric repeats. The new putative promoters used in this study ranged in size from 1585 base pairs (bp) to 2599 bp (Supplementary Table 1; Supplementary Figure 2). Each putative promoter consisted of a 5’UTR of 55 bp to 373 bp and an intron of 874 bp to 1114 bp. The *S. italica* putative promoter contained the longest 5’UTR of 373 bp, which was interrupted by a second intron of 48 bp.

**Table 1.**
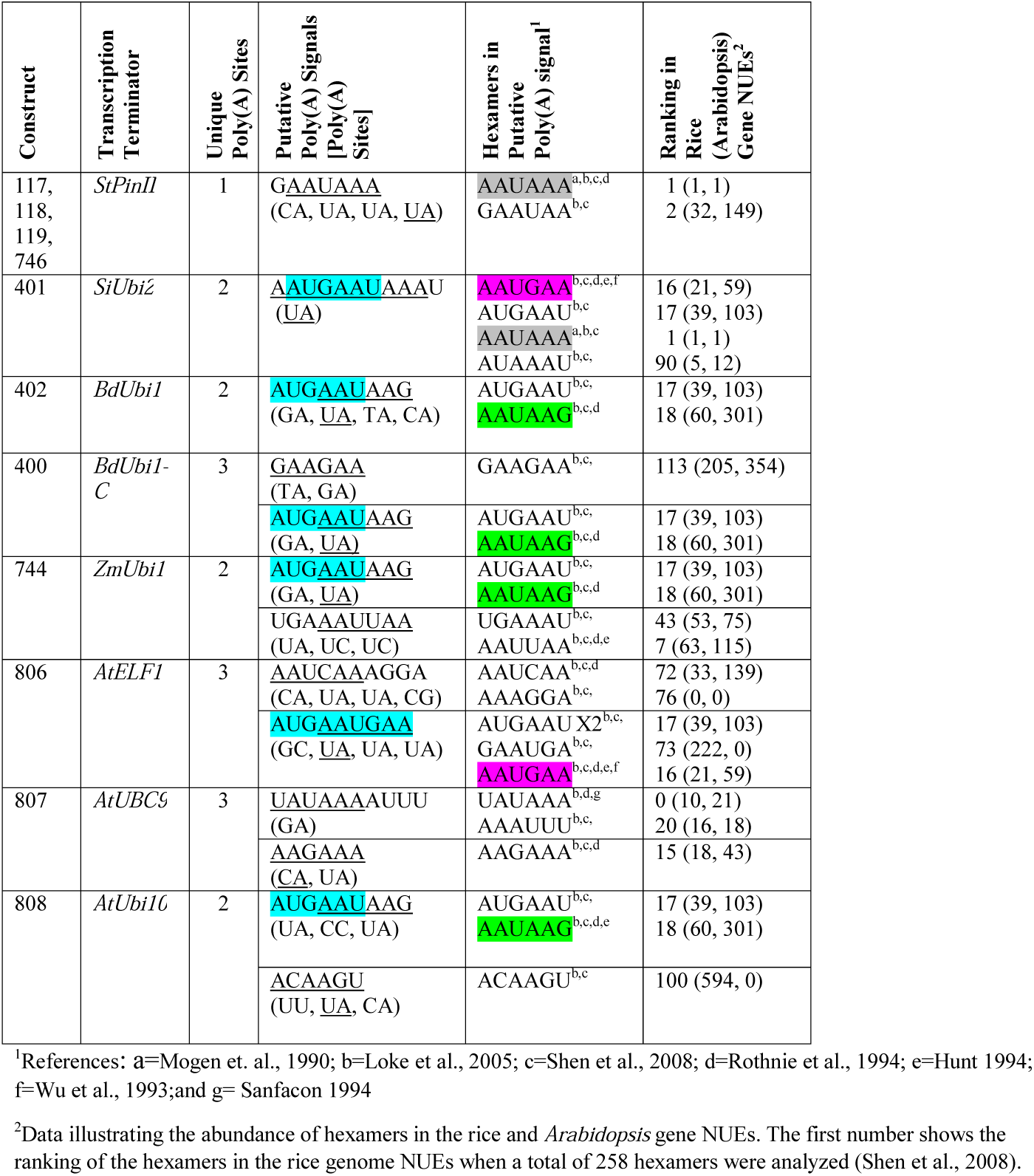
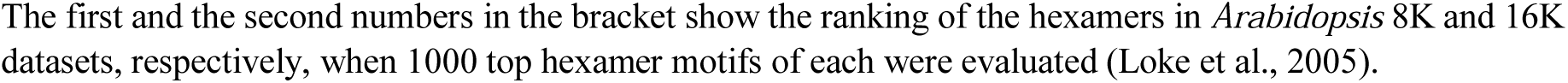
Analysis of putative poly(A) signals (PPSGs) in various transcription terminators (TTs). PPSGs (underlined), AUGAAU, an overlapping top-ranking hexamer of the rice and *Arabidopsis* near upstream elements (NUEs) is highlighted in turquoise, two-letter poly(A) sites are shown in brackets in the line underneath (major sites underlined) in distinct TTs. Hexamers of rice and *Arabidopsis* gene NUEs (either PPSGs or overlapping hexamers) are marked with superscripts (either b or c or both in column 5) showing the reference where they were reported. Gray, magenta and green highlighted sequences are PPSGs that occurred more than once in the TTs.

Putative promoter sequences of *BdUbi1, SiUbi2*, and *BdUbi1-C* were aligned with that of the *ZmUbi1*, and limited sequence homology was identified within the REs (Supplementary Figure 3). Similarly, putative TT sequences derived from the four UBQ genes were aligned (Supplementary Figure 4) and they revealed 33 - 47% homologies between any two sequences. The highest sequence homology of 47% was identified between the *BdUbi1* and *BdUbi1-C* TTs. All REs were deployed in assembling chimeric constructs (Figure 1A and 1B) to evaluate gene expression in transgenic maize.

**Figure 3.**
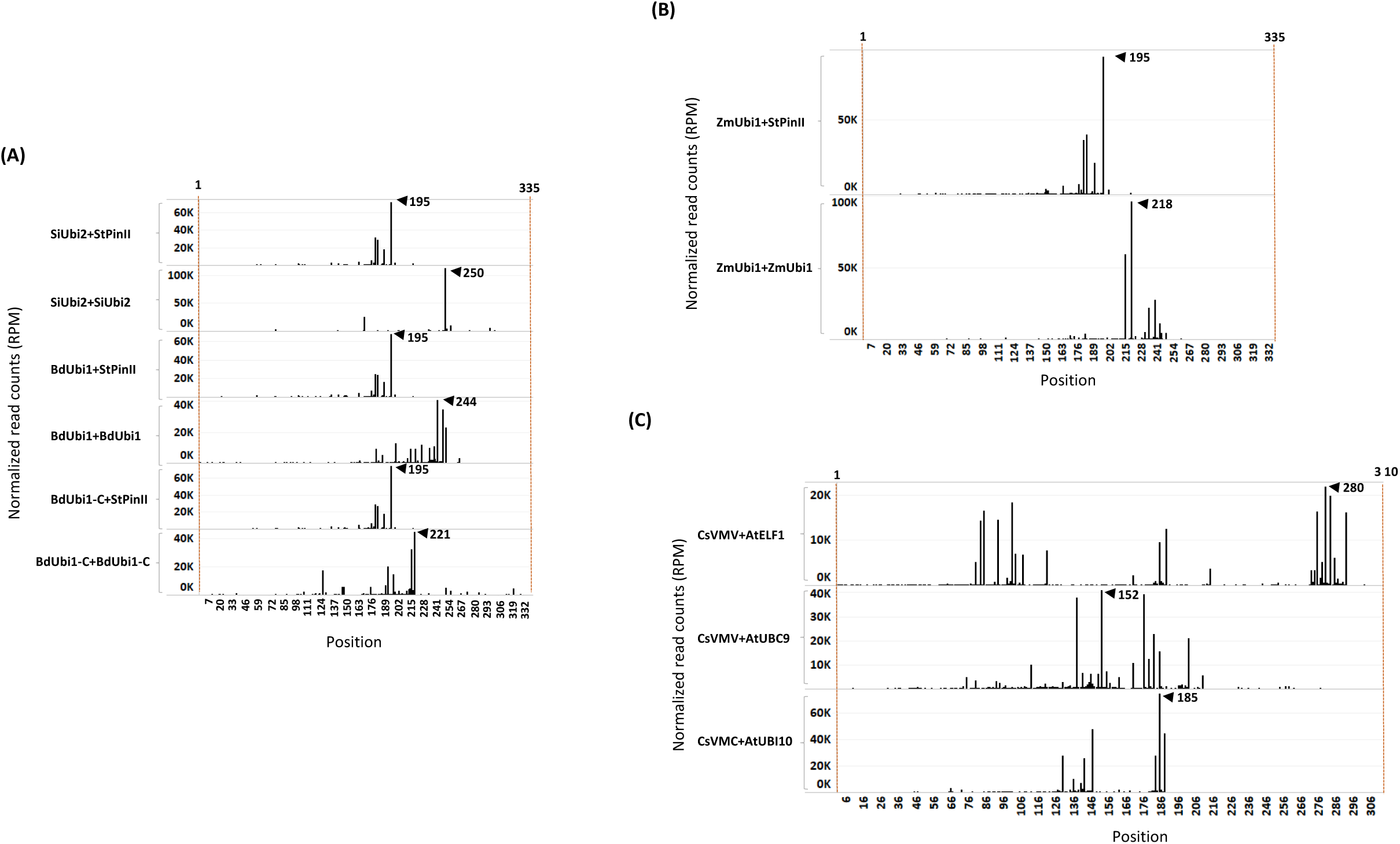
3’ RACE sequence mapping profiles of mRNA extracted from V6 maize leaves at the T_1_ generation. mRNA poly(A) sites and its abundance reflected in individual peaks are shown for distinct TTs. On the X-axis, the position 1 starts at the last base of the reporter gene. The Y-axis indicates the normalized reads number per million (RPM) of the 3’ RACE deep sequencing read end (after trimming poly (A) mapping for each position). The peak of poly(A) site representing mRNA abundance is determined using a cut-off of 10,000 RPM. For each construct, two individual samples were sequenced, and similar results were observed. The representative data is shown here in the sequence mapping profiles of up to 335 nt. (A) Comparison of constructs containing the *StPinII* TT (*SiUbi2+PinII*; *BdUbi1-C+PinII, BdUbi1+PinII*), *SiUbi2* TT (*SiUbi2+SiUbi2*), *BdUbi1* TT (*BdUbi1+BdUbi1*) and *BdUbi1-C* TT (*BdUbi1-C+BdUbi1-C*). (B) Comparison of *StPinII* TT (*ZmUbi1+StPinII*) and *ZmUbi1 TT* (*ZmUbi1+ZmUbi1*). (C) Comparison of *AtELF1* TT (CsVMV+*AtELF1*), *AtUBC9* TT (CsVMV+*AtUBC9*) and *AtUbi10* TT (*ZmUbi1+AtUbi10*). The number indicated by the black triangle represents the position of the highest peak (highest mRNA abundance) for that construct.

**Figure 4.**
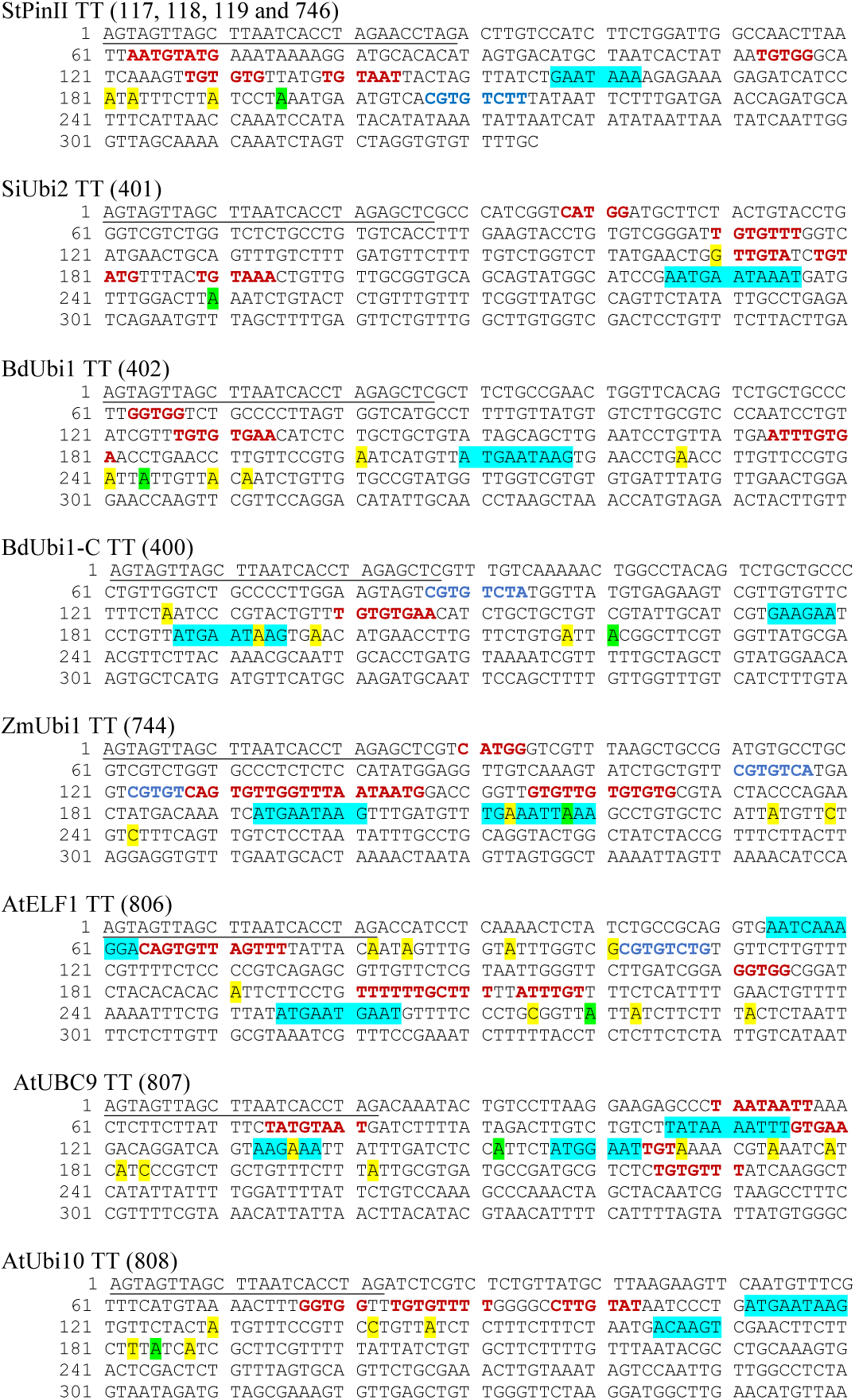
DNA sequence of distinct TTs deployed in constructs of transgenic maize. The first bp in the sequence is the last bp of the reporter gene and coordinate 1 of the diagrams in Figure 3. The underlined sequences were present between the reporter gene coding sequence and the TTs. The turquoise, green and yellow highlighted sequences depict PPSGs and overlapping hexamers as shown in Table 1, major poly(A) sites representing most abundant mRNA as shown in Figure 3, and non-major poly(A) sites as shown in Figure 3, respectively. The letters in red and blue font are putative FUE motifs and mRNA stability motifs (e.g., CGTGTCTT of *StPinII*), respectively, that were previously described (Supplementary Table 2). If two motifs overlapped, only one of the motifs is shown.

### 3.2. Analysis of Gene Expression

All chimeric constructs were evaluated by analyzing mRNA and protein expression of the reporter genes, *yfp* and *cry34Ab1*, in T_1_ maize leaves at V6. To compare the strengths of the TTs, each of the four UBQ promoter expression was compared in pairs of chimeric constructs when the reporter gene was terminated either by the *StPinII* TT or the native TT derived from the same gene as the promoter (Figure 1). Higher mRNA expression was observed in the *SiUbi2*+*SiUbi2* (promoter+TT; this order is the same throughout this manuscript) and *ZmUbi1*+*ZmUbi1* constructs containing the native combinations of REs as compared to the *SiUbi2*+*StPinII* and *ZmUbi1*+*StPinII* constructs, respectively in transgenic maize (Fig. 2A and 2B). Both of these differences were up to 3-fold higher and statistically significant. The mRNA expression appeared higher in the native combinations of *BdUbi1+BdUbi1* and *BdUbi1*-*C*+*BdUbi1-C* constructs as compared to their respective *BdUbi1+StPinII* and *BdUbi1-C+StPinII* combinations, but the differences were not statistically significant. The mRNA expression observed for the positive control constructs, *ZmUbi+StPinII*, containing two different reporters (Fig. 2A and 2B) was similar, ∼ 5 units each, indicating that mRNA expression was consistent despite utilizing different reporter genes. The protein expression in all four constructs containing the native combinations of UBQ REs was 2- to ∼3.4-fold higher than that of the respective UBQ promoter and *StPinII* TT combinations (Figures 2C and 2D). All four of these differences were statistically significant. The highest difference was observed between *ZmUbi1+ZmUbi1* and *ZmUbi1+StPinII* constructs (Figure 2D). The protein expression in *ZmUbi1+StPinII* construct containing two different reporters (Figure 2C and 2D) could not be compared due to different ELISA assays used for the two reporter genes.

To further understand gene expression by varying TTs, the viral CsVMV promoter was paired with three distinct TTs derived from the *A. thaliana* genes, *AtELF1, AtUBC9* and *AtUbi10* in chimeric constructs (Figure 1B). Based on the analysis of the maize plants transgenic for one of these three constructs, CsVMV+*AtUbi10* produced the highest *cry34Ab1* mRNA expression, ∼1.8 fold higher than CsVMV+*AtELF1* (Figure 2B). This difference was statistically significant. The mRNA expression differences between CsVMV+*AtUbi10* and CsVMV*+AtUBC9* were not statistically different. *Cry34Ab1* protein expression levels of CsVMV+*AtUbi10* was highest, >4-fold higher than CsVMV+*AtELF1*, and >2-fold higher than CsVMV*+AtUBC9* (Figure 2D). These differences were statistically significant. Interestingly, the protein expression levels of CsVMV+*AtUbi10* and *ZmUbi1+ZmUbi1* were similar, indicating that these combinations were equally effective in expressing the reporter gene. Taken together, these results demonstrated that varying TTs could have a significant impact on gene expression regulation in transgenic maize.

### 3.3. Small RNA analysis for gene silencing

Transcriptional and post-transcriptional regulation of gene expression mediated by small RNA (sRNA or siRNA) of 21-24 nt in length has been previously demonstrated by Matzke and Mosher (2014). Presence of antisense siRNA is a signature of siRNA-mediated gene silencing. To investigate whether mRNA and protein expression differences observed among and within the construct pairs were due to sRNA-mediated transgene silencing, sRNA sequencing was performed in T_1_ maize leaves transformed with chimeric constructs (Figure 1). The siRNA analyses of plants individually transformed with twelve constructs did not reveal any antisense siRNA (Supplementary Figure 5). Background levels of sense siRNA were observed in a few cases, which could be the result of mRNA degradation as reported previously (Sidorenko et al., 2017).

### 3.4. Analysis of polyadenylation sites in transcription terminators

NGS following 3’ RACE analysis was performed to investigate the number of poly(A) sites within the TTs, and which of the ensuing mRNA isomers contributed to the majority of protein expression. Alignment of the 3’ RACE-sequencing reads (RPM) with the corresponding gene TT identified poly(A) sites and their relative quantities in the V6 leaves of T_1_ maize plants (Figure 3). Groups of poly(A) sites present within the 10-35 nt window were considered as one unique site. This is based on the observations that canonical AAUAAA polyadenylation motif is consistently present within 10-30 nt from the poly(A) site in *Arabidopsis* (Loke et al., 2005) and 10-35 nt in rice (Shen et al., 2008). Polyadenylation machinery produces redundant cleavage-driven poly(A) sites in order to dismantle the machinery from mRNA (Shen et al., 2008). Constructs containing the StPinII TT, e.g., *SiUbi2+StPinII, BdUbi1+StPinII, BdUbi1-C+StPinII* and *ZmUbi1+StPinII*, each yielded four poly(A) sites (or one unique poly(A) site) within the 35 nt window consistently (Figures 3A and 3B; Figure 4). The majority mRNA polyadenylated at 195 nt, and this position is shown as a major peak for these constructs. These results illustrated that the *StPinII* TT was consistent in polyadenylating the pre-mRNA at pre-defined sites and was not influenced by any of the four promoters or two reporters present on the same construct in transgenic maize (Figure 1A and 1B).

Similar to *SiUbi2+StPinII, BdUbi1+StPinII, BdUbi1-C+StPinII*, constructs bearing the native combinations of the UBQ REs often produced two or more poly(A) sites within the 35 nt span, and these sites were considered as one unique poly(A) site (Figure 1A). Accordingly, *SiUbi2+SiUbi2, BdUbi1+BdUbi1* and *BdUbi1*-*C*+*BdUbi1-C* produced two, two (six total), and three (five total) unique poly(A) sites, respectively (Figure 3A; Figure 4). Among these poly(A) sites, the major poly(A) sites were located at 250 nt, 244 nt and, 221 nt for constructs *SiUbi2+SiUbi2, BdUbi1+BdUbi1* and *BdUbi1*-*C*+*BdUbi1-C*, respectively. The native combination of the *ZmUbi1+ZmUbi1* REs exhibited 2 unique (5 total) poly(A) sites (Figure 3B, Figure 4). The major site was located at 218 nt. All poly(A) sites in the UBQ TTs were located within the 260 nt from the end of the reporter gene (or from nt 1 shown in Figures 3 and 4).

The CsVMV+*AtELF1*, CsVMV+*UBC9 and* CsVMV+*Ubi10* chimeric constructs, identified different patterns of poly(A) sites (Figure 3C). Construct CsVMV+*AtELF1* and CsVMV+*UBC9* exhibited three unique poly(A) sites each (totals of nine and eight, respectively), while construct CsVMV+*AtUbi10* exhibited two unique poly(A) sites (six total). The most abundant poly(A) site is shown in Figure 3C. All poly(A) sites were located within 300 nt from the end of the reporter gene.

### 3.5. Sequence analysis of polyadenylation sites and flanking cis elements in transcription terminators

Although the plant consensus signal for the poly(A) site is YA (CA or UA), alternative poly(A) sites have been identified in TTs (Cui et al., 2003; Loke et al., 2005). Of the 45 poly(A) sites identified in eight TTs, 66.7% cleaved pre-mRNA either at CA or UA (Figure 4). Other alternative poly(A) sites were, GA (15.6%), UC and CU (8.9%), GG and GC (6.7%), and UU (2.2%).

Sequence analysis of the near upstream element (NUE) regions of the eight TTs, located 10 to 35 nt distal to the poly(A) sites, identified canonical AAUAAA and AAUAAA-like poly(A) signals in the TTs. These are referred to as putative poly(A) signals (PPSG) as none have been validated through conventional mutagenesis studies or other methods in maize. Of these PPSGs, only two TTs contained the canonical poly(A) signals, the *StPinII* and *SiUbi2* (Figure 4). The *StPinII* and *SiUbi2* TTs produced a major poly(A) site 32 nt and 15 nt downstream of this signal, respectively (Figure 4). The AAUAAA-like PPSGs were identified in the TTs and they included AAUAAG, AAUGAA, AAUCAA, AAUUAA, UAUAAA and AAGAAA. These PPSGs were verified previously either through conventional mutagenesis or linker scanning studies in plants [reviewed in (Wu et al., 1993; Hunt, 1994; Rothnie et al., 1994; Sanfaçon, 1994; Wu et al., 1995)]. AAUAAA-like PPSGs were identified in five TTs, *SiUbi2* (overlapped with AAUAAA), *ZmUbi1, BdUbi1, BdUbi1-C*, and *AtELF1*, and they were present within 26 nt upstream of the major poly(A) sites (Figure 4). Two AAUAAA-like PPSGs, AAUAAG and AAUGAA were most frequent in these five TTs (Table 1). The *AtUbi10* TT also contained AAUAAG, nevertheless, it generated only three non-major poly(A) sites. The major poly(A) site was present 65 nt downstream of this PPSG. A variant of AAUGAA, AAUGAC, was identified 19 nt upstream of the major poly(A) site, but this motif was absent in rice and *Arabidopsis* NUEs. Overlapping with the AAUGAA motif, an ACAAGU motif was located 15 nt upstream of the major poly(A) site and this could be the PPSG for the *AtUbi10* TT. This identification is based on the bioinformatics data available for the *Arabidopsis* and rice gene NUE motifs (Loke et al., 2005; Shen et al., 2008), the A-richness of the motif and its proximity to the major poly(A) site (Table 1). For the *AtUBC9* TT, the AAGAAA motif located 14 nt upstream of the major poly(A) site was identified as a PPSG. For the non-major poly(A) sites present in the TTs, either AAUAAA-like PPSGs were identified, including AAUUAA in *ZmUbi1*, AAUCAA in *AtELF1*, UAUAAA in *AtUBC9* (Table 1) or the poly(A) signal was ambiguous due to sequence variability in plants (Hunt, 1994; Rothnie et al., 1994). The exception was the GAAGAA motif identified as a PPSG for the *BdUbi1-C* TT based on its A-richness, proximity to the poly(A) sites, and its abundance in *Arabidopsis* and rice gene NUEs (Table 1). Overall, PPSGs were identified for ∼72% of the unique poly(A) sites (Figure 4; Table 1).

To find out whether the PPSGs occurred alone in the TTs or overlapped with other hexamers, all TTs were analyzed with 20 and 50 top-ranking hexamers (Loke et al., 2005; Shen et al., 2008) of the NUEs of rice and *Arabidopsis* genes respectively. Accordingly, one specific hexamer, AUGAAU, overlapped with PPSGs in six of the eight TTs analyzed, except for *AtUBC9* and *StPinII* (Table 1; Figure 4). In five of the six such overlaps, a major poly(A) peak was observed within 10 nt-26 nt downstream of this composite motif. The final size of such overlapped motifs was 9-11 nt. Of these, AUGAAUAAG was observed in the *ZmUbi1* TT and AAUGAAUAAAU in the *SiUbi2* TT (Table 1). Interestingly, the 9 nt motif was identical in the *ZmUbi1, BdUbi1, BdUbi1-C* and *AtUbi10* TTs, and it was an overlap of AUGAAU and AAUAAG PPSGs. The 11 nt motif of the *SiUbi2* terminator was a composite of four hexamer motifs, the canonical AAUAAA and AAUGAA, and two hexamers (AUGAAU and AUAAAU) that ranked high in rice and *Arabidopsis* gene NUEs (Table 1). Furthermore, all four monocot TTs (*ZmUbi1, BdUbi1, BdUbi1-C* and *SiUbi2*) consisted of only one copy of the AUGAAU and overlapped with PPSG. However, the *AtELF1* TT consisted of two copies of AUGAAU present in a direct repeat, AUGAAUGAAU and the AAUGAA PPSG was embedded in it. Other composite motifs included the overlap of GAAUAA with canonical AAUAAA in the *StPinII* TT, resulting in a heptamer motif, GAAUAAA (Table 1). This heptamer motif alone ranked 16^th^ in the *Arabidopsis* gene NUEs (Loke et al., 2005), implying the significance of this overlap.

In *ZmUbi1, BdUbi1-C*, and *AtUBC9* TTs, at least two PPSGs were identified that were spatially separated. The upstream PPSG putatively generated a poly(A) site within the downstream PPSG. In *ZmUbi1* TT (ZmUbi1+ZmUbi1), two distinct PPSGs, AAUAAG and AAUUAA, were identified occurring 12 nt apart (Fig. 4). The upstream PPSG generated two poly(A) sites within the downstream PPSG corresponding to the highest 213 nt and 218 nt poly(A) peaks (Figure 3B). A similar phenomenon was observed in the *BdUbi1-C* (*BdUbi1-C+BdUbi1-C*) and *AtUBC9* (CsVMV+*AtUBC9*) TTs, whereby a poly(A) site was observed within the downstream PPSGs, AAUAAG and AAGAAA, respectively. Despite that, the downstream PPSGs appeared functional as they created a major poly(A) peak downstream of the motifs (Figures 3A and 3B, Figure 4). These results could be examples of alternative polyadenylation (APA) and require further investigation.

GC-rich motifs reportedly increase mRNA stability when located downstream of the poly(A) sites. Such a GC-rich motif, CGUGUCUU, was identified in the *StPinII* TT 12 nt downstream of the major poly(A) site (An et al., 1989). Similar motifs were identified in the FUEs of three TTs. The *ZmUbi1* TT contained two such motifs, CGUGUC<u>A</u>U and CGUGUC<u>AG</u> with 1-2 nt variations as underlined. The *AtELF1* and *BdUbi1-*C TTs each had one GC-rich motif, CGUGUCU<u>G</u> and CGUGUCU<u>A</u>, respectively (Supplementary Table 2). The first 6 nt of these two GC-rich motifs were conserved as compared to that in the *StPinII* TT. Whether these sequences have any significance to increase the stability of mRNA requires further investigation (Hunt, 1994).

## Discussion

Typically, promoters and TTs in chimeric constructs have been paired from diverse sources, including *Agrobacterium* and viruses, and across dicotyledonous and monocotyledonous plant species (An et al., 1989; Richter et al., 2000; Diamos and Mason, 2018). Nevertheless, there have been few examples where promoters and TTs were paired from the same gene (Hiwasa-Tanase et al., 2011). In this study, seven novel UBQ gene REs (promoters and TTs) from *SiUbi2, BdUbi1, BdUbi1-C*, and *ZmUbi1* were identified. Constructs were assembled carrying the native combinations of these REs, and individual promoter and *StPinII* TT combinations, to study transgene expression in maize. To best mimic the native expression of the REs, the REs were cloned using seamless cloning, which replaced the UBQ coding sequence with reporter genes to avoid sequence modification (Zhu et al., 2007; Kumar et al., 2017a). Like the *ZmUbi1* promoter, the *BdUbi1, BdUbi1-C*, and *SiUbi2* promoters contained introns that have been shown to enhance gene expression (Rose, 2008). Despite that, expression driven by the *ZmUbi1* promoter was the highest observed of the four UBQ promoters, irrespective of the TT with which it was paired. The varied expression levels observed with the other promoters are valuable for transgene expression in maize. Depending on the trait genes to be expressed, a wide range of expression is required in multi-gene constructs. There have been multiple studies reporting the identification of UBQ gene promoters from diverse plant species and characterization of their expression in transgenic plants [summarized in (Lu et al., 2008; Mann et al., 2011)]. However, pairing of the UBQ promoters with their native TTs added further flexibility in designing constructs in commercially important maize.

mRNA stability, localization, and translational efficiency are associated with the sequences in the 3’ UTR regions of the TTs (Proudfoot, 2004; Gilmartin, 2005). However, when TTs are used in a heterologous transgenic system, it is crucial to confirm their efficacy to ensure desired level of expression. Compared to the control constructs containing the *StPinII* TTs, increased expression of reporter genes was observed when UBQ promoters were paired with their respective gene TTs. These results indicated that the UBQ TTs possibly improved one or more of the following functions: mRNA polyadenylation, stability, translocation and translation (Sachs and Wahle, 1993; Li and Hunt, 1997; Hiwasa-Tanase et al., 2011), independent of the promoter controlling transcription. Increased expression observed from the CsVMV promoter when paired with the *Arabidopsis* TTs further supports this notion.

Another potential hypothesis for increased expression is improved transcription termination from the UBQ TTs as compared to that from the *StPinII* TT control. The UBQ promoters and UBQ TTs can provide enhanced cross-talk for efficient RNA Polymerase II machinery function (Mapendano et al., 2010; Andersen et al., 2012). Formation of un-polyadenylated or aberrant transcripts were reported when weaker terminators were paired with high expressing viral promoters (Luo and Chen, 2007). These aberrant transcripts often lead to gene silencing through the RNA-dependent RNA polymerase pathway (Matzke and Mosher, 2014). The sRNA analysis did not reveal the presence of antisense siRNA in the T_1_ progenies derived from any of the constructs used in this study (Supplementary Figure 5), indicating that the levels of aberrant transcripts, if generated, were not sufficient to trigger the RDR6-mediated gene silencing pathway. These findings suggest that the lower levels of transgene expression observed in multiple constructs containing *StPinII, AtELF1*, or *AtUBC9* TTs, were not due to transgene silencing. Similar to these results, the *StPinII* TT is reported to provide multiple-fold higher accumulation of target gene products than that of the *nos* TT (An et al., 1989; Richter et al., 2000; Diamos and Mason, 2018) and transgene expression with *StPinII* TT was at similar levels to that observed with other soybean TTs (Xing et al., 2010). Accordingly, the constructs containing the *StPinII* TT did not exhibit any gene silencing.

Polyadenylation signals that generate multiple poly(A) sites within the 10 nt to 35 nt stretch of pre-mRNA are a characteristic of the plant TTs (Dean et al., 1986; Hunt, 1994; Loke et al., 2005; Shen et al., 2008). Among the TTs studied in this report, up to three unique poly(A) sites were observed. All poly(A) sites were confined within 300 nt from the end of the reporter gene. Similarly, as many as 19 unique poly(A) sites are reported in TTs of rice genes and the average length of the 3’ UTR is 289 nt in rice (Shen et al., 2008). This 3’ UTR length is close to the ∼300 nt length observed in this study. The unique sites could represent APA because maize leaf contains multiple tissue types. Each tissue type could be processing the poly(A) sites uniquely due to tissue-specific gene regulation. A specific phenomenon was observed in the *ZmUbi1* TTs in this study, whereby the upstream PPSG presumably created a major poly(A) site within the downstream PPSG located 12 nt apart. This step might have allowed the generation of shorter transcripts as compared to the mRNA isoforms generated by the downstream PPSG. It is feasible that the two PPSGs are active in different cells/tissues due to APA, as a maize leaf is comprised of multiple tissue-types. Shorter mRNAs are transcribed in actively dividing cells in the early stages of development (and perhaps in rapidly diving cells of V6 maize leaves used in these experiments), whereas longer mRNA are synthesized in the later stages of development regulated by the motifs (e.g., miRNA binding sites, RNA binding protein factors, and AU-rich elements(Barreau et al., 2005; Fabian et al., 2010), controlling gene expression (Sandberg et al., 2008; Mayr and Bartel, 2009; Di Giammartino et al., 2011). Therefore, shorter RNAs synthesize more proteins. In *BdUbi1-C* and *AtUBC9* TTs, the upstream PPSG generated a non-major poly(A) site within the downstream PPSG but it was the downstream PPSG that created the major poly(A) site within 30 nt downstream. The length of the 3’ UTR can affect stability. transport, localization and translational properties of mRNA. Deep sequencing of the *Arabidopsis* transcriptomes identified ∼10% of the transcriptome harboring APA that may be specific to developmental stages and tissues responding to environmental stresses (Shen et al., 2011). Nevertheless, the observations made in the present study for APA warrant additional studies to confirm cell/tissue-specific expression during maize leaf development.

An intriguing feature of the majority of TTs examined herein was the presence of composite PPSGs consisting of the canonical AAUAAA or AAUAAA-like motifs, and their overlaps with either other PPSGs or top-ranking hexamers of *Arabidopsis* and rice gene NUEs resulting in 9 to 11 nt motifs [Table 1; (Loke et al., 2005; Shen et al., 2008)]. Longer poly(A) signals have been reported previously in the TT of pea *1,5 ribulose biphosphate carboxylase* gene (*rbcS-E9*) site 1, such as AAUGGAAAUGGA, which is a duplication of the AAUGGA motif, and was verified through conventional mutagenesis studies (Hunt, 1994). Although *rbcS-E9* is a highly expressed gene in plants, this hexamer motif is not the top-ranking hexamer of the *Arabidopsis* gene NUEs, and ranked 244 of the 1000 hexamer patterns analyzed (Loke et al., 2005; Shen et al., 2008). This result indicated that TTs of high expressing genes do not necessarily contain the top-ranking hexamers in their poly(A) signals. Another longer poly(A) signal for the TT of wheat histone H3 gene, AAUGGAAAUG, was verified in mutagenesis studies (Ohtsubo and Iwabuchi, 1994). This sequence is identical to the first 10 nt of the *rbcS-E9* site 1 poly(A) signal. The last six nt of this sequence, GAAAUG, also occurs in the rice (ranked 23 of the 258 hexamers identified) and *Arabidopsis* (ranked 187 in 1000 patterns analyzed) gene NUEs. The *ocs* site 2 polyadenylation signal is a 10 nt motif, AAUGAAUAUA, identified through linker scanning studies (Macdonald et al., 1991). The first 8 nt of this motif, AAUGAAUAU, is identical to that of the *SiUbi2* PPSG, AAUGAAUAAAU (Table 1). It is noticeable that both *ocs* site 2 and *SiUbi2* PPSGs contained the prevalent AUGAAU motif, which is a top-ranking hexamer of the rice and *Arabidopsis* gene NUEs. Composite motifs could strengthen the poly(A) signal. This hypothesis was supported when clustering of multiple motifs of a repeat, TTTGTA, and its variants in the FUE of the CaMV 35S poly(A) signal strengthened this signal during transfection of *Nicotiana plumbaginifolia* leaf protoplasts (Rothnie et al., 1994). This strengthening of the signal resulted in induction of 3’ end processing at a heterologous polyadenylation site in the cryptic *nos* locus.

Among the multiple hexamer motifs identified that overlapped with PPSG in the TTs (Table 1), the AUGAAU motif overlapped in six of the eight identified PPSGs (Table 1). The AUGAAU motif ranked 17^th^ in the TTs of rice gene NUEs and occurs 2331 times (0.0012 frequency) as compared to 4177 times (0.0015 frequency) for the canonical AAUAAA polyadenylation motif (Shen et al., 2008). Similarly, in *Arabidopsis*, the AUGAAU motif ranked 39^th^ [Table 1; (Loke et al., 2005)]. These numbers showed a significant presence of the AUGAAU motif in the NUEs of the rice and *Arabidopsis* TTs and its putative role in 3’ end processing. After inspection of 15 different poly(A) signals in the petunia chlorophyll a/b binding TT, a similar AUGAAA motif was identified as a PPSG (Dean et al., 1986). The significance of the overlap of AUGAAU with PPSGs is unknown at this time and warrants further investigation.

As shown here, the AUGAAU motif was prevalent in the TTs of high expressing UBQ genes, especially that of the monocotyledonous plants. All four monocotyledonous TTs carried the AUGAAU motif in their PPSGs, which generated a major poly(A) site 14 to 26 nt downstream. The PPSGs of the *ZmUbi1, BdUbi1* and *BdUbi1-C* TTs carried the identical motif, AUGAAUAAG, yet their mRNA abundance profiles were distinctly different (Figure 3). As NUE polyadenylation signals and cleavage sites (CSs) control the utilization of poly(A) sites in a TT, the FUE motifs located upstream of the NUEs can enhance the efficiency of polyadenylation at one or more poly(A) sites (Rothnie, 1996; Li and Hunt, 1997). Therefore, the sequence, number, and spatial arrangement of all three *cis* elements, FUE, NUE and CS, of a polyadenylation signal are crucial in determining the mRNA poly(A) sites and their abundance in TTs. Accordingly, the FUE motifs of the *ZmUbi1, BdUbi1*, and *BdUbi1-C* TTs varied in sequence, number, and their relative locations (Supplementary Table 2; Figure 4). The distinctly different poly(A) patterns and mRNA abundance profiles observed with these three TTs could be due to identified FUE motif differences.

The *AtUbi10* TT also contained the composite PPSG motif, AUGAAUAAG, identical to three of the four monocotyledonous UBQ TTs. Unlike the monocotyledonous TTs, the *AtUbi10* TT PPSG produced only three non-major poly(A) sites. The major poly(A) site was located 65 nt downstream to this PPSG. Similar polyadenylation profile of mRNA abundance and isoforms was observed in the native genome of *Arabidopsis* for the AtUbi10 TT [<u>TAIR10</u>(Duc et al., 2013)]. These results indicated that the composite AUGAAUAAG PPSG may function differently due to spatial arrangement of the *cis* elements in the *AtUbi10* TT and their interaction in transgenic maize (Hunt, 1994).

In summary, the present study provides insights into construct design and composition for single- and multi-gene constructs for transgene expression in maize. These findings may apply to other plant species. Novel promoters and TTs were identified and characterized for transgene expression in maize. Compared to control *StPinII* TT, higher transcript abundance and protein accumulations were observed with the novel TTs when used in combinations with their native promoters. Small RNA analysis did not reveal any evidence of gene silencing from any of the constructs. The sequence analysis of these TTs revealed the presence of composite PPSGs consisting of the canonical AAUAAA or AAUAAA-like NUE motifs, and their overlaps with multiple top-ranking hexamers of *Arabidopsis* and rice gene NUEs, specifically AUGAAU. The AUGAAU was present in six of the eight TTs studied. Further insights into the functional significance of these overlaps in mRNA 3’ end processing could be gathered through analysis of transgene expression from constructs carrying synthetic TTs (modified) or *via* conventional mutagenesis studies. In addition, the present studies utilized traditional transformation methods and were insightful despite complexities caused by genome context of random transgene insertion. Targeted, site-specific insertion is now possible to reduce the potential impact of genome location on transgene insertion, and such studies may improve precision of transgene expression (Ainley et al., 2013; Kumar et al., 2016; Anand et al., 2019; Betts et al., 2019; Gao et al., 2020).Taken together, the identification of unique promoters and TTs sequences generating high transgene expression will be very useful for creating molecular transgene stacks for future biotech crops.

## Author Contributions

PW, SK and MG conceived and designed the experiments, supervised the execution of research, and analyzed the data. TW contributed to regulatory element strategy and discussions. RM generated the NGS data and JZ analyzed the NGS data. PW and SK assembled the Figures and MG wrote the manuscript. All authors critically reviewed and approved the manuscript.

## Acknowledgement

We are grateful to Andrew Asberry, Wei Chen, Shavell Gorman, Avani Manduri (summer intern), Greg Schulenberg and Andrew Worden for nucleic acid and protein analyses; to Maize Transformation Team for transformation of constructs in maize; and to Jamie Lutz for the care of plants in the greenhouse. The authors also thank Dr. Rodrigo Sarria and Michelle Smith for their support during this research. We acknowledge Tracey Fisher for the critical review of the manuscript.

## Figure legend

**Supplementary Figure 1**. Protein sequence alignment of *Z. mays* (ZM) *Ubi1, S. italica Ubi2, B. distachyon Ubi1* and *B. distacyon Ubi1-C* UBQ genes. The highlighted amino acids represent consensus among two (turquoise) or all four (yellow) sequences.

**Supplementary Figure 2**. Nucleotide sequences of the *BdUbi1-C* (2A), *BdUbi1* (2B) and *SiUbi2* (2C) genes depicting putative promoters (underlined sequences distal of ATG contain TSS and upstream sequences), 5’ UTR (upper case), intron (gray highlighted), protein coding sequences (italicized), and TT regions (downstream of TAA, a translational stop codon).

**Supplementary Figure 3:** Nucleic acid alignment of the maize *polyubiquitin1* (ZM Ubi1) gene promoter sequence with *B. distachyon* (*Ubi1* and *Ubi1-C*) and *S. italica* (*Ubi2*) UBQ promoter sequences using Vector NTI software. The promoter sequences include the “total length” shown in the Supplementary Table 2. The highlighted nucleotides represent consensus among two (green), three (turquoise), or all four (yellow) sequences. All four sequence consensus is also highlighted with red font.

**Supplementary Figure 4:** Nucleic acid alignment of the maize *polyubiquitin1* (*ZmUbi1*) gene TT sequence with putative TTs derived from the *S. italica* (Si*Ubi2*) *and B. distachyon* (*BdUbi1* and *BdUbi1-C*) UBQ genes. The alignment was carried out using Clustal W 2.1 software. The non-highlighted nucleotides represent the nucleotide homologies between one or more sequences. Whereas gray highlighted nucleotides represent differences between one or more sequences. The entire consensus sequence is gray highlighted indicating sequence differences at every single nucleotide among the four UBQ TTs.

**Supplementary Figure 5.** siRNA analyses of distinct constructs. Sequencing of sRNA was performed as described in Materials and Methods, and the transgene-derived siRNA accumulation is shown here. For each construct, two individual events (as shown) were selected for whole-genome sRNA sequencing. The siRNA was mapped to each component of the transgene cassettes, including promoter (P), protein coding sequence, and TT with the differentiation of sense and antisense strand (denoted on top). The sRNA accumulation shown here is read counts normalized to the total reads derived from the corresponding library and depicted as stacked bars with colors signifying siRNA sizes (i.e., 21-24 nt). The comparison of siRNA accumulation is presented here for constructs *ZmUbi1+StPinII* (746) and *ZmUbi1+ZmUbi1* (A); CsVMV*+AtELF1*, CsVMV+*AtUBC9* and CsVMV+*AtUbi10* (B); *SiUbi1+StPinII, SiUbi1+SiUbi1, BdUbi1+StPinII, BdUbi1+BdUbi1, BdUbi1-C+StPinII* and *BdUbi1-C+BdUbi1-C* (C).

**Supplementary Table 1:** Length of the putative promoters and TTs for the UBQ genes.

**Supplementary Table 2**. Putative FUE motifs identified in various TTs that have been reported previously to function as 3’ end processing signals. A subset of the motifs observed in the rice and *Arabidopsis* gene FUEs and putative stabilization signals are also depicted. The underlined sequences show nt variations in previously reported sequences as shown.

