## Supplementary-Figure 3 for "Transcription terminator-mediated enhancement in transgene expression in maize: preponderance of the AUGAAU motif overlapping with poly(A) signals"

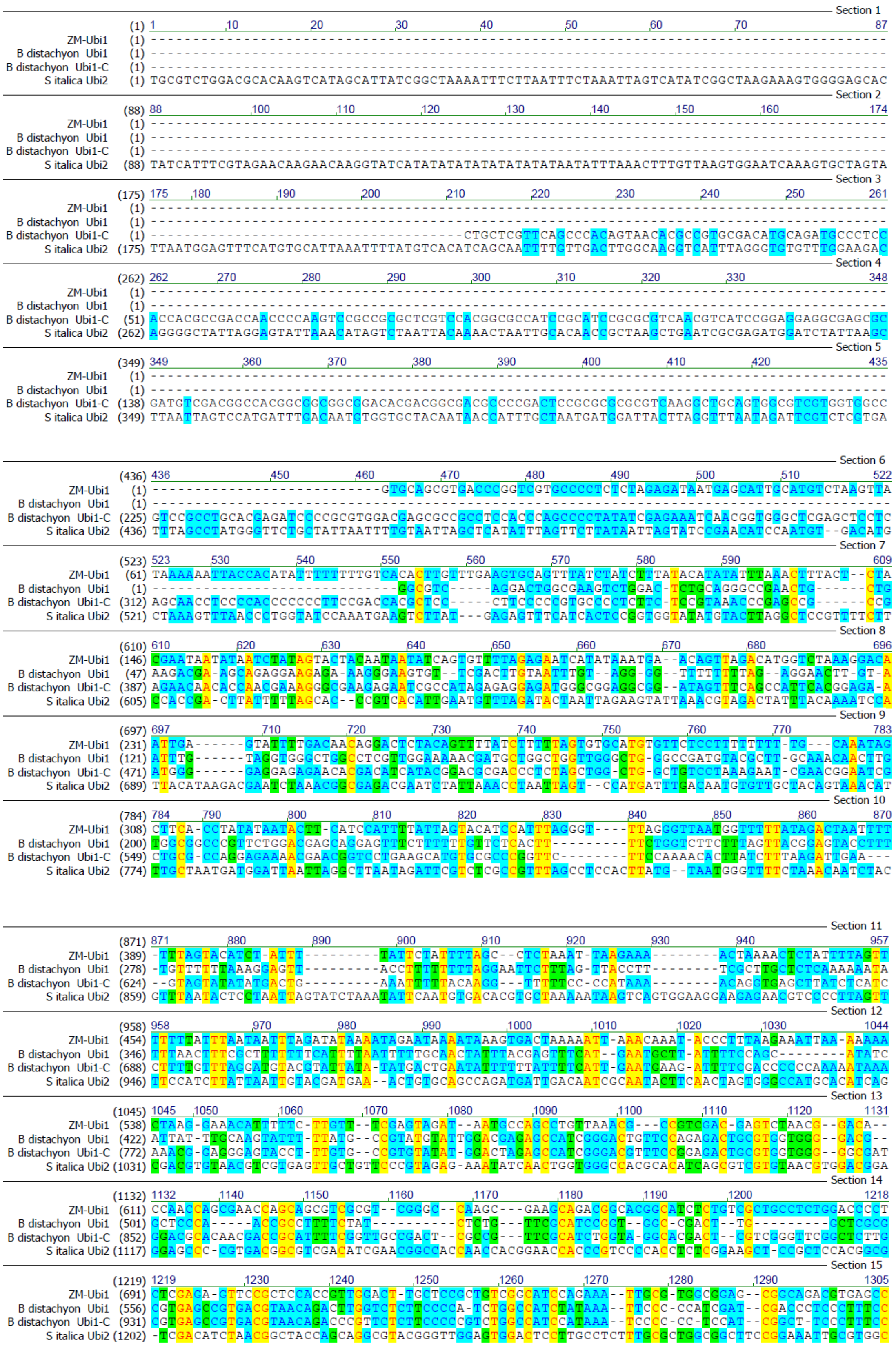

```
Section 16
(1306)                 1306  1320  1330  1340  1350  1360  1370  1380  1392
ZM-Ubi1            (771)  GGCACGGCAGGCGGCCTCCTCCTCCTCTCAC-GGCACGGCAGCTACGGGGGATTCCTTTCCCACCGCTCCTTCGCTTTCCCTTCCTC
B distachyon Ubi1  (637)  C-CAATCCAGCACCCCCGATCCCGATCGAAAATTCTCCGCAACAGCAAGCGAT-----------CGATCTAGCGAATCCCCGTCAAG
B distachyon Ubi1-C (1011) T-CAATCCAGCACCCT-GATTCCGATCGAAAAGTCCCCGCAAGAGCAAGCGAC-----------CGATCTCGTGAATCTCCGTCAAG
S italica Ubi2    (1288)  G-GAGACGAGGCGGGCTCGTC----TCACAC-GGCACGGAAGACGTCACGGGTTCCTTCCCCACCTCTCCT---CTTCCCCACCGCC

Section 17
(1393)                 1393  1400  1410  1420  1430  1440  1450  1460  1479
ZM-Ubi1            (857)  GCCCGCCGTAATAAATAG---ACACCC----CCTCCACACCCTCTTTCCCCAACCTCGTGTTGTTCGGAGCGCAC-ACACACA-CAA
B distachyon Ubi1  (712)  G----------TATGTAG---CCTCTCGATTCCTCCTCAGCCCTGCCCTCGATT-TGGTGTACG-CGTTGAG-------------AT
B distachyon Ubi1-C (1085) G----------TATGCAG---CCTCGC--TTCCTCCTCGCTACCGTT-TCAATTCTGGAGTAGGTCGTAGAGGAT-AC------CAT
S italica Ubi2    (1366)  A----------TAAATAGCCGACCCCCTCGCCTTTCTCCCCAATCTCATCTCGTCTCGTGTTGTTCGGAGCACACCACCCGCCCCAA

Section 18
(1480)                 1480  1490  1500  1510  1520  1530  1540  1550  1566
ZM-Ubi1            (935)  CCAGATCTCCCCAAATCCACC-CGTCGGCACCTCCGCTTCAAGGTACGCCGCTCGTCCTCCCCCCCCCCCCCTCTCTACCTTCTCT
B distachyon Ubi1  (771)  GATGAT------------------------CTCGTAGAT----GTC---------------TAGATGACACCATGTCGATTTGAAAT
B distachyon Ubi1-C (1149) GTTGATTTGACAGAGGGAGTAG-ATTAGATACTTGTAGATCGAAGTGCGGATGTTCCATGGTAGATGATACCATGTTGATTTCGATT
S italica Ubi2    (1443)  ATCGTTCTTCCCGCAAGCCTCGGCGATCCTTCACCCGCTTCAAGGTACGGCGATCGTCTT--CCTCCTCTAGATCGGCGTGATCTGC

Section 19
(1567)                 1567  1580  1590  1600  1610  1620  1630  1640  1653
ZM-Ubi1           (1021)  AGATCGGCGTTCCGGTCCATGGTTAGGGCCCGGTAGTTCTACTTCTGTTCATGTTTGTGTTAGATCCG---TGTTTGTGTTAGATCC
B distachyon Ubi1  (815)  AGATCAGA--------------T-----CCGTGTAGATCGA--TGAGCTCCTGTG-------------------TACCTGTGGATTC
B distachyon Ubi1-C (1235) AGATCGGA--------------TTAAATCTTTGTAGATCGA--AGTGCGCATGTTCCA-TGAATTGCC---TGTTACCAGTAGATTC
S italica Ubi2    (1528)  AAGTAGTTGATTTGGTAGATGGTTAGGATCTGTGCACTGAA---GAAATCATGTT------AGATCCGCGATGTTTCTGTTCGTAGA

Section 20
(1654)                 1654  1660  1670  1680  1690  1700  1710  1720  1730  1740
ZM-Ubi1           (1105)  GTGCTGCTAGCG------TTCGTACA---CGGATGCGA---CCTGTACGTCA-GACACGTTCTGATTGCTAACTTGCCAGTGTTTCT
B distachyon Ubi1  (862)  AAGTTATTTT-------------------------------CG-----------------------------CATGCTATTGT----
B distachyon Ubi1-C (1302) AAGTTTTTCTGT------GTTATAGAGGTGGGATCTAC---TCGTTGAGATG-ATTAGCTCCTAGAGGACACCATGCCGTTTTGGAA
S italica Ubi2    (1606)  TGGCTGGGAGGTGGAATTTTTGTGTAGATCTGATATGTTCTCCTGTTTATCTTGTCACGCTCCTGC-GATTTGTGGGGATTTTAGGT

Section 21
(1741)                 1741  1750  1760  1770  1780  1790  1800  1810  1827
ZM-Ubi1           (1179)  CTTTG----GGGAATCCTGGGATGGCT-CTAGCCGTTCCGCAGACGGGATCGATTTCATGATTT------TTTTTGTTTCGTTGCA-
B distachyon Ubi1  (885)  -------------------------------TGTGATC----------------------------------------TACTAG---
B distachyon Ubi1-C (1379) AATAG----ATCAGAACCGTG-TAGAT-CGATGTGAGCATGTGTTCCTGTAGATCCAAGTTCTT------TCGCATGTTACTAGTTG
S italica Ubi2    (1692)  CGTTGATCTGGGAATCGTGGGGTTGCTTCTAGGCTGTTCGTAGATGAGGTCGTTCTCACGGTTTACTGGATCATTGCCTAGTAGATC

Section 22
(1828)                 1828  1840  1850  1860  1870  1880  1890  1900  1914
ZM-Ubi1           (1254)  ----TAGGGTTTGGT--TTGCCCTT--TTCCTTTATTTCAATATATG---CCGTGCACTTGT-TTGTCGGGTCATCTTTTCATGCTT
B distachyon Ubi1  (898)  ------ATCTAGTGT--GTG---TA--TTCTATGCTATCGATTTCT----CCGTGTAGATTT-CACTCGATTACTGTTACTGTGGCT
B distachyon Ubi1-C (1454) ----TGATCTATTGT--TTG---TG--TAATACGCTCTCGATCTAT----CCGTGTAGATTT-CACTCGATTACTGTTACTGTGGCT
S italica Ubi2    (1779)  AGCTCGGGCTTTCGTCTTTGTATATGGTGCCCATACTTGCATCTATGATCTGGTTCCGTGGTGTTACCTAGGTTTCTGCGCCTGATT

Section 23
(1915)                 1915  1920  1930  1940  1950  1960  1970  1980  1990  2001
ZM-Ubi1           (1329)  TTTTTTGTCTTGGTTGTGATGATGTGGTCTG-GTTGGGCGGTCGTTCTA----GATCGGAGTAGAATT--CTGTT-TCAAACTACCT
B distachyon Ubi1  (967)  TGATCGGCCATAGATGTTG-GTTAAGGTTTG-ATCGGTTAGTGTTTGAACCT-GCGTGGATATCTAGC--ATCCA-TCTATTATCGT
B distachyon Ubi1-C (1525) TGATCGTTCATAGTTGTTC-GTTA-GGTTTG-ATCGAACAGTGTCTGAACCT-AATTGGATATGTATT--CTTGA-TCTATCAACGT
S italica Ubi2    (1866)  CGTCCGATCGATTTTGTTAGCATGTGGTAAACGTTTGGTCATGGTCTGATTTAGATTAGAGTCGAATAGGATGATCTCGATCTAGCT

Section 24
(2002)                 2002  2010  2020  2030  2040  2050  2060  2070  2088
ZM-Ubi1           (1408)  GGTGGATTTATTAATTTTGGATCTGTATGTGTGTGCCATACATATTCATAGTTACGAATTGAAGATGATGGA-TGGAAATATCGATC
B distachyon Ubi1 (1048)  GTAGGTTTCGAACAAACAAGCACTATTATTGTACTGATGGTTCGTCTATGGTTG-GTTTGACCGTTTAGTGTTGAACGAGCCTTC
B distachyon Ubi1-C (1605) GTAGGTTTC----AGTCATG---TATTTATGTACTCC--CTCCGTCCCAAATTA-ACT--GACGTGGATTTTGTATAAGAATCTATA
S italica Ubi2    (1953)  CTTGGATTAATATGCATGTGTCACCAA-TCTGTTCC---G-TGGTTAAGATGATGAATCTATGCTTAGTTAATGGGTGTAGATATA

Section 25
(2089)                 2089  2100  2110  2120  2130  2140  2150  2160  2175
ZM-Ubi1           (1494)  TAGGATAGGTATACATGTTGATGCGGGTTTACTGATGCATATACAGAGATGCTTTTGTTCGCTTGGTTG-TGATGATGTGGTGTG
B distachyon Ubi1 (1134)  TG---TATT--TGTTTATTGCTGTCCAGTGA--TGTACCATGTTCGTTGAGTGTCGGATTATACTAATTAT-TGTTGAT------TG
B distachyon Ubi1-C (1680) CA---AATCCATGTCAGTTAATTCGGGATGG--AGTACCATATTCAAT---------AATTTGTTTATTGC-TGTCCAC------TT
S italica Ubi2    (2035)  TATGCTGCTGTTCCTCAATGATGCCGTAGCTTTACCTGAGCAGCATGGATCCTCC-TGTTACTTAGGTAGATGCACATGCTTATAG

Section 26
(2176)                 2176  2190  2200  2210  2220  2230  2240  2250  2262
ZM-Ubi1           (1580)  GTTGGCGGTCGTTCATTCGTTCTAGATCGGAGTAGAATACTGTTTCAAACTACCTGGTGTATTTATTAATTTTGGAACTGTATGTG
B distachyon Ubi1 (1207)  ATAATCTTGTAGTTTGCTTTTCCTAATTTAT-TTATCGTAGTCCTG--ATTTGCCTCAGCTGTGCCTCACCCGTGCGATGGTCAATC
B distachyon Ubi1-C (1746) ATGTACCATATGTTTGTTGTTCCTCATGTGGATTCTACTAATTATC--ATT-GATT--GGTGATCTTCTATTTTGCTA--GT-----
S italica Ubi2    (2121)  ATCAAGATATGTACTGCTACTGTTGGAATTCTTTAGTATACCTGATGATCATCCATGCTCTTGTTACTTGTTTTGGTATACTTGGAT

Section 27
(2263)                 2263  2270  2280  2290  2300  2310  2320  2330  2349
ZM-Ubi1           (1667)  TGTGTCATACATCTTCATAGTTACGAGTTTA------AGATGGATGGAAATATC----GATCTAG--------GATAGGTATACATG
B distachyon Ubi1 (1291)  AACTTGTTAGCCCAATCTGCTTAATCATGTA------CATTTGTTGTTAGAATCAGA-GATCAAG--------CCA---ATTAGCTA
B distachyon Ubi1-C (1821) ---TTCCTAGCTCAATCTGGTTATTCATGTA------GATGTGTTGTTGAAATCGGA-GACCATG--------CTTGTTATTAGATA
S italica Ubi2    (2208)  GATGGCATGCTGCTGCTTTTTGTTGATTTGAGCCCATCCATATCTGCATATGTCACATGATTAAGATGATTACGCTGTTTCTGTATG

Section 28
(2350)                 2350  2360  2370  2380  2390  2400  2410  2420  2436
ZM-Ubi1           (1736)  TTGATGTGGGTTTTA-CTGATGCATATACA---TGATGGC-ATATGCAGCA-TCTATTCA---TAT-----GCTCTAACCT--TGAG
B distachyon Ubi1 (1360)  TCT-TATTGCTTAT--CTGTTCCATGTTC----TGATCG----ATGTAACA-G-TCTACACT-TTT-----GCTCTGTGCT--ACTT
B distachyon Ubi1-C (1890) GTT-TATTGCTTAT--CAGTTTCATGTTC----TGGTTG----ATGCAACA-CATATTCATG-TTC-----GCTATCTGGT--TGCT
S italica Ubi2    (2295)  ATGCCATAGCTTTTATGTGAGCAACATGCATCCTCCTGGTTATATGCATTAATAGATGGAAGATATCTATTGCTACAATTTGATGAT

Section 29
(2437)                 2437  2450  2460  2470  2480  2490  2500  2510  2523
ZM-Ubi1           (1807)  TACCTATCTATTATAATAAACAAGTATGTTTTATAATTATTTGATCTTGATATACTTGGATGATGGCATA-TGCAGCAGCT--AT-
B distachyon Ubi1 (1426)  GATTAAAACATTCTGACTTAAATTCATGATTGGAAGTT--TCAGATCTG---ATTGTTGCCTTA---CTTG-ACTAATATCT--ATT
B distachyon Ubi1-C (1957) GCTTGATATTCTCTGATTTACATTCATTATAAGAATATATTCTGCTCTG---GTTGTTGCTTCT---CATG-ACTT-TACCT--A--
S italica Ubi2    (2382)  TATTTGTACATACGATGATCAAGCATGCTCTTCA--TACTTGTTGAT---ATACTTGGATAATGAAATGCTGCTGCACGTTCATT

Section 30
(2524)                 2524  2530  2540  2550  2560  2570  2580  2590  2600  2610
ZM-Ubi1           (1890)  -ATGTGGATTTTTTAGCCCTGCCTTCATACGCTATTTAT-TTGCTTGGTACTGTTTCTTTTGTCGATGCTCACC-----CTGTTGT
B distachyon Ubi1 (1502)  CATGTGACACCTCTCTGTCTTGGTAACTTACCGC---TGT-TTGTTTGTAATTTCTGACTATGCAG---------------------
B distachyon Ubi1-C (2032) ------------CTCGGTA--GGTGACTTACC-----TTT-TGGTTTACAATTGTCAACTATGCAG---------------------
S italica Ubi2    (2464)  CTATAGCACTAATGATGTGATGAACACGCACGACCTGTTTGTGGCATCTGTTTGAATGTGTTGTTGCTGTTCACTAGAGACTGTTTT
```

```
                 (2611) 2611     2620       2630       2640       2650      2660
          ZM-Ubi1 (1970) TTGGTGTTACTTCTGCAG--------------------------------
  B distachyon  Ubi1 (1564) --------------------------------------------------
B distachyon  Ubi1-C (2078) --------------------------------------------------
     S italica Ubi2 (2551) ATTAACCTACTGCTAGATACTTACCCTTCTGTCTGTTTATTCTTTTGCAG
```
