## Supplementary figures and images for "Transcription terminator-mediated enhancement in transgene expression in maize: preponderance of the AUGAAU motif overlapping with poly(A) signals"

### Supplementary-Figure 5

#### Supplementary Figure 5

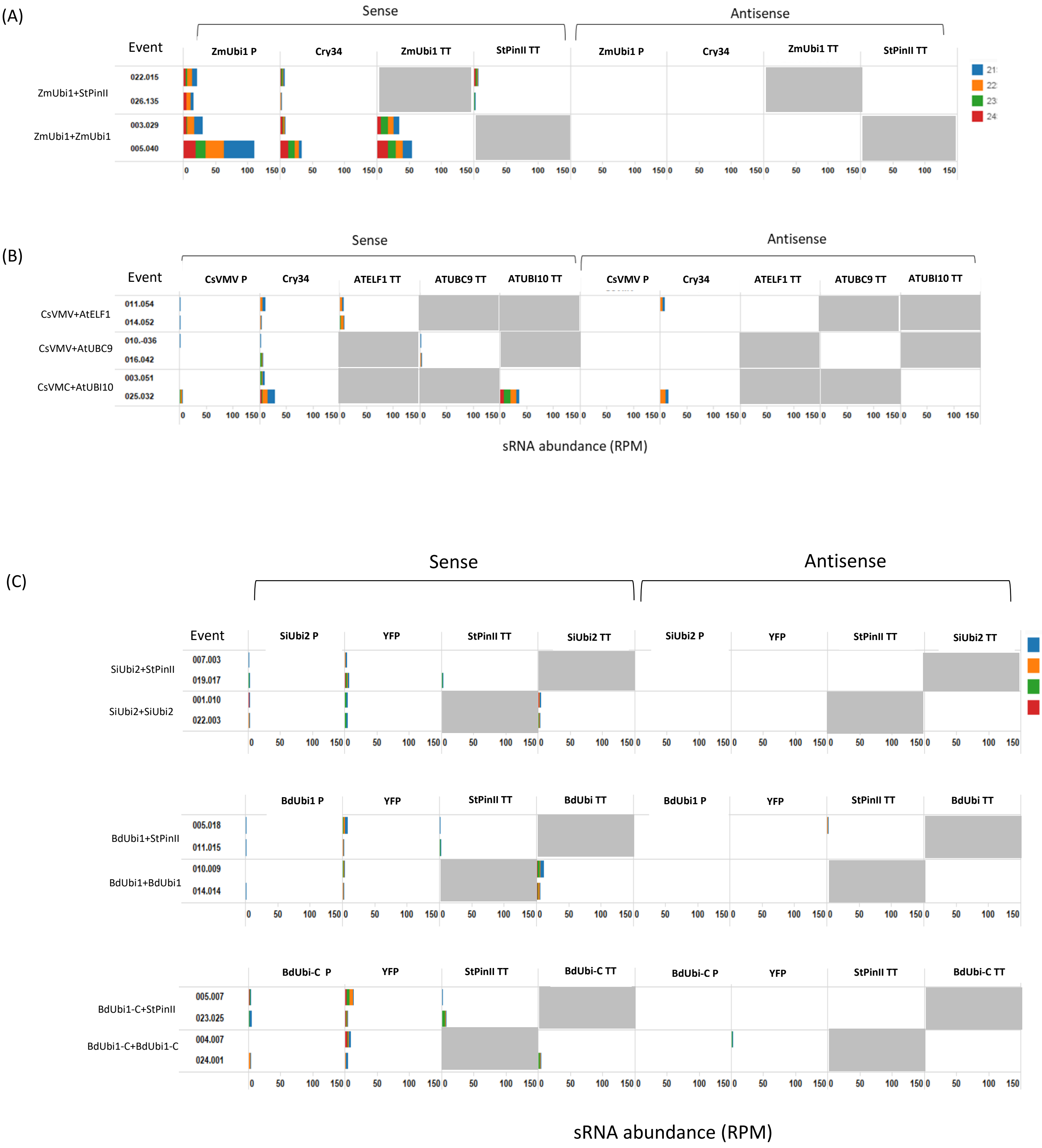
