## Supplementary-Figure 4 for "Transcription terminator-mediated enhancement in transgene expression in maize: preponderance of the AUGAAU motif overlapping with poly(A) signals"

```
Consensus     GSYCATGGKTCTTNNAARCTGSYNCWSWGYCTGCKKCSYCTGGTG---GYCTGYCYCT--    55
ZmUbil TT     -GTCATGGGTCGTTTAAGCTGCCGATGTGCCTGCGTCGTCTGGTG---CCCTCTCTCCAT    56
SiUbi2 TT     GCCCATCGGTCATGGATGCTTCTACTGTACCTGGGTCGTCTGGTCTCTGCCTGTGTCACC    60
BdUbil TT     ------GCTTCTGCCGAACTGGTTCACAGTCTGCTGCCCTTGGTG---GTCTGCCCCT--    49
BdUbil-C TT   ------GTTTGTCAAAAACTGGCCTACAGTCTGCTGCCCCTGTTG---GTCTGCCCCT--    49

Consensus     -TGGAAGTAGTCRTGTC----TTTTGTTATGTGN--N--TGTTGTGTG--NNYCAWKYCC   104
ZmUbil TT     ATGGAGGTTGTCAAAGT----ATCTGCTGTTCG------TGTCATGAG----TCGTGTCA   102
SiUbi2 TT     TTTGAAGTACCTGTGTCGGGATTGTGTTTGGTCATGAACTGCAGTTTGTCTTTGATGTTC   120
BdUbil TT     -T---AGTGGTCATGCC----TTTTGTTATGTG------TCTTGCGTC----CCAATCCT    91
BdUbil-C TT   -TGGAAGTAGTCGTGTC----TATGGTTATGTGAGAAGTCGTTGTGTTCTTTCTAATCCC   104

Consensus     GTATTGTTTGTGTGAWSAWCTGSTKSTGTYGTRT-TGCATMCTGAANANTCCTGTTAYGA   163
ZmUbil TT     GTGTTGGTTTAATAATGGACCGGTTGTGTTGTGTGTGCGTACTACCCAGAACTATGACAA   162
SiUbi2 TT     TTTTGTCTGGTCTTATGAACTGGTTGTATC-TGTATGTTTACTGTAAACTGTTGTTGCGG   179
BdUbil TT     GTATCGTTTGTGTGAACATCT-CTGCTGCTGTAT-AGCAGCTTGAA---TCCTGTTATGA   146
BdUbil-C TT   GTACTGTTTGTGTGAACATCTGCTGCTGTCGTAT-TGCATCGTGAAGAATCCTGTTATGA   163

Consensus     ATCAGTGAACNTGAACCTTGWTSTGTGAATNNNNTTANGACTTAG-TSATTATGCTCTTT   222
ZmUbil TT     ATC-ATGAAT---AAGTTTGATGTTTGAAAT---TAAAGCCTGTGCTCATTATGTTCTGT   215
SiUbi2 TT     TGCAGCAGTATGGCATCCGAATGAATAAATGATGTTTGGACTTAAATCTGTACTCTGTTT   239
BdUbil TT     ATTTGTGAACCTGAACCTTGTTCCGTGAATCATGTTATGAATAAG-TGAACCTGAACCTT   205
BdUbil-C TT   ATAAGTGAACATGAACCTTGTTCTGTGA------TTACGGCTTCG-TGGTTATGCGAA--   214

Consensus     GTTT-CGKTTTTAYCATTACAATTKCNYSTGAGGTMWGRTTGTTT-GYGTTTGMTTTMTG   280
ZmUbil TT     CTTT-CAGTTGTCTCCTAATATTTGCCTGCAGGTACTGGCTATCT-ACCGTTTCTTACTT   273
SiUbi2 TT     GTTTTCGGTTATGCCAGTTCTATATTGCCTGAGATCAGAATGTTTAGCTTTTGAGTTCTG   299
BdUbil TT     GTTC-CGTGATTATTGTTACAATCTGTTGTGCCGTATGGTTGGTC-GTGTGTGATTTATG   263
BdUbil-C TT   -----CGTTCTTACAAACGCAATTGCACCTGATGTAAAATCGTTT-TTGCTAGCTGTATG   268

Consensus     TTGARCTKGTKGACGACGYTCRTTCCAGAAGTTAKTSN-CAACCTTWGYTRRAACATCCW   339
ZmUbil TT     AGGAGGTGTTTGAATGCACTAAAACTAATAGTTAGTGGCTAAAATTAGTTAAAACATCCA   333
SiUbi2 TT     TTTGGCTTGTGGTCGACTCCTGTTTCTTACTTGAGGCGTAACTCTGTTCTGGCAAACTCA   359
BdUbil TT     TTGAACTGGAGAACCAAGTTCGTTCCAGGACATATTG--CAACCTAAGCTAAACCAT-GT   320
BdUbil-C TT   GAACAAGTGCTCATGATGTTCATGCAAGATGCAATTC--CAGCTTTTGTTGGTTTGTCAT   326

Consensus     AATACY--ACTGNTTNTAGTAGAMMTAAAANGTNNTTYTKRTTAATTNGTTWGTTKTGWK   397
ZmUbil TT     AACACC--ATAGCTAATAGTTGAACTATTAGCTATTTTTGGAAAATTAGTTAATAGTGAG   391
SiUbi2 TT     AATGTCTAACTGAATGTTTTAGGACTTAATTGTTGGACAGATTAACGTGTTTGGTTTGTT   419
BdUbil TT     AGAACT--ACTTGTTCTGGGAGACATAAAACGTCATTTTTATGCATTCGTAACATTTAAG   378
BdUbil-C TT   CTTTGT--ACTGTGCTTACCGCACATAAAGATTGCATCTTGCTTATTGCTTTGTTGCTTT   384
```

```
Consensus    GNTRCTATN-TRTTCGCWWGGYCTTTYACCTAATCAWTCTT-GAGANCAGATANCTCTTY   455
ZmUbil TT    GTAGTTATT-TGTTAGCTAGCTAATTCAACTAACAATTTTTAGCCAACTAACAATTAGTT   450
SiUbi2 TT    TCTAGATTGTGATTCGGAAGGCTTGTTAGTTGTGGAATCAAGGAGAGCAGCTAGGTCTGT   479
BdUbil TT    CATACTACAATAATTGTATTGTCCTTTTCCTACTCATCCTT-GAAACCATATGCCTCTTC   437
BdUbil-C TT  GGTGCT---CGTCCGCTTCTCCTTGCACCTTATCAAACCT-TTGTTTAGAT-TCTCTTC   438

Consensus    TCAGAGCATT-TANWNRCTCTNAGCTTTATAACAACATGGCTCTAYCTGMARYYTCATRA   514
ZmUbil TT    TCAGTGCATT-CAAACACCCC---CTTAATGTTAACGTGGTTCTATCTACCGTCTCCTAA   506
SiUbi2 TT    GCAGAACGTTATTTTGGATTTAAGCCTTCTCAGATTATGCCATTACTCTAAACCTAATGA   539
BdUbil TT    TCAGCGCCTC-TACATGCAGTGTGCTC-AGAACAAACAGGCCCTGCCAGCTGCTTTTCAA   495
BdUbil-C TT  TTATAGCACT-TGGTAACTCTCAGCTTTACAACGCCAGTACTGTTTCTGAAATTTCATGA   497

Consensus    TWT-AT--TTAAYTGATAGATGTAGTASTAGKM-TWTGGCWTCTGTTGGTGACTATTYGA   570
ZmUbil TT    TAT-ATGGTTGATTGTTCGGTTTGTTGCTATGCTATTGGGTTCTGATTGCTGCTAGTTCT   565
SiUbi2 TT    TATCATATTTCACTCGGGGATGTTGGAGTAGTC-TTTTCTTTCTCCTGCAGACAAAATGA   598
BdUbil TT    TTT--T--CCAATTAATAACCACAATAGTCGGACTATGGCATCTGTGGGTGACTATGCAA   551
BdUbil-C TT  CTG-AT--AAAGCTGATAGATGGAGTACTAATA-TATGACATCTTTCCATAAATGTTCGG   553

Consensus    KTNTGNWKCYRTGAGKKCTC-TRRTATTG-TCCNATTTATATGATGAATTTACCTGRTAA   628
ZmUbil TT    TGCTGAATCCAGAAGTTCTCGTAGTATAGCTCAGATTCATATTATTTATTTGAGTGATAA   625
SiUbi2 TT    TTTTGCTTTCGTGTGTGTACATGATTTTG-TGCAACTGTTGCAACAACTGAAGTAGACAA   657
BdUbil TT    G-ATGTTGCTGTCAGGTCTC-TGAAACTTTTCCCATGTATCTGTTGAAATTACCCAGTAA   609
BdUbil-C TT  GTGCAGAGATATGGAGGCCC-CAGGA---TCCTATTTACAGGATGAACCTACCTGGGCC   608

Consensus    GTTNTANCCWCGAYATAAGCATGCAAG----TTTGRNNTTTTYWACAWGGAYAWRTTATT   684
ZmUbil TT    GT--GATCCAGGTTATTACTATGTTAG----CTAGGTTTTTTTACAAGGATAAATTATC   679
SiUbi2 TT    GTTTTGACCTCACCAGAAGAATGAAAAAGATTTTGGAATTTGTTACATCGACAAACCATT   717
BdUbil TT    ATTCATGCCTCTATTTAATCTGGCATG----GTTGA--TTTTCAAACAGAATGTGTTTTT   663
BdUbil-C TT  GCTGTACGCATGACATCCGCGAGCAAG----TCTGAGGTTCTCAATGTACACATGAAATT   664

Consensus    KTTTNTTGTNGCATYTRTATTGGCWGAWNRTTGCMTAGATGCWGWGTCATGATATTTN-A   743
ZmUbil TT    TGTGATCATAATTCTTATGAAAGCTTTATGTTTCCTGGAGGCAGTGGCATGCAATGC--A   737
SiUbi2 TT    GTAACTTGGCCCATCAGAATGCACAGAAGAGCGGCTACAAATTGACATGCGTTGCAAACT   777
BdUbil TT    TTTTGTTCTGGAAGCTATATTGGTAAATAAATACAAAGCTGGAGTGTGATTATATTT--C   721
BdUbil-C TT  GATTTTTGCTGCGTTTGGCTTGGCTGATCGTTGCATTTGTTCTGATTCATCAGAGTTAAA   724

Consensus    TAACAGAWATTTAAGCAMATNTCAGCTGATG-YAKCYACNACTG-RGTANATRTATGGTT   801
ZmUbil TT    TGACAGCAACTTGATCACA--CCAGCTGAGG-TAGATAC------GGTA---ACAAGGTT   785
SiUbi2 TT    TTGCAATAGTTGATGCACATGTTTGCCATTGCCTGCCAGTCTTAGGAAAAGTGTGTGGTT   837
BdUbil TT    CAACAGATATTCAAGAAAATCTCAGTTGATT-TATTTACTACTGTAGTATATATATATAT   780
BdUbil-C TT  TAACGGATATATCAGCAAATATCCGCAG----CATCCAC-ACCG-ACCACACGTCCGGTT   778
```

```
Consensus     C-WTAAAKYTGACCWCCCATGCTTTRAWCAACAYGCGWGCACAYYGYTCAGATTCTT---   857
ZmUbi1 TT     C-TTAAATCTGTTCACCAAATCATTGGAGAACACACATACACATTCTTGCCAGTCTTGGT   844
SiUbi2 TT     CGAGAAATCTAAGCATATGTGCTCTGCTCACATTGCGTGGAACCCACACAGCTTTGT---   894
BdUbi1 TT     C-TTACAGTTGACTTCTCATATTTCAAACGACATGTGAGCACATTGTTCAGTTTCTT---   836
BdUbi1-C TT   A-ACAGAGTCCCCCTGCCTTGCTTTAATTATTACG-GAGTACTCCGCTATTAATCCT---   833

Consensus     TAGATATGTTTCGWGNCMAAANGTCAAWTCTKGCATTNTTCACYCCTAGTRAACANTAAC   917
ZmUbi1 TT     TAGAGAAATTTCATGACAAAATGCCAAAGCTGTCTTGACTCTTCACTTTTGGCCATGAGT   904
SiUbi2 TT     CACACTCTTGTCCACTCCAGAAGTCATTCCTGGCGCTGTTTACCCCTGGTAAAAGGTAAC   954
BdUbi1 TT     AGGATGTGTTGTGTGCTCAAAGGTGTAATTTTGCATTCTGCCCTCCGAGTAAACACTACA   896
BdUbi1-C TT   TAGATATGTTTCGAAGGAACTCAAACCTTCCTCCATCTGCAAATCTCAGTGCTTCAAAAC   893

Consensus     CGTAATTTTHTVADTGVHABNTBCATTHGDTTACAANNVDCAACAGCAAATCDAAVABCT   977
ZmUbi1 TT     CGTGAC------------------------------------------------------   964
SiUbi2 TT     CGAAAACTTCTCAAGGCTGTACCCAAAACTGGAAGGAAATTTGGAGGAAATCTTTGCTTT  1014
BdUbi1 TT     CGTATTTTTTTGAGTGGCAG-TGCATTTGATTACAA--GGCAACAACAA--CAAAAACCT   951
BdUbi1-C TT   TGGAATTAGATAATTGAAACCTTCATTCGGTTGCAATTCACAACTGCAAATTGAACAGCA   953

Consensus     HTGTCAABTTAHHCTTCNNNTTMRMGRYTSCMMSGATMRKTTKRMMTGAWCYATGWWMSR  1037
ZmUbi1 TT     ------------------------------------------------------------  1024
SiUbi2 TT     TGATCGGCTCACTCTTTC------------------------------------------  1074
BdUbi1 TT     ATGGCAAGATATCCTTC---TTAGAGGCTGCCAGGATCATTTTGACTGAACTATGTAAGG  1008
BdUbi1-C TT   CTGTCAATTTCAATTTCGGGTTCACGATTCCACCGATAGGTTGACATGATCCATGATCCA  1013

Consensus     CYSAWKRWAMRRC                                                1050
ZmUbi1 TT     -------------                                                 910
SiUbi2 TT     -------------                                                1032
BdUbi1 TT     CTGAAGAAAAGG-                                                1020
BdUbi1-C TT   CCCATTGTACAAC                                                1026
```
