## Supplementary-Figure 1 for "Transcription terminator-mediated enhancement in transgene expression in maize: preponderance of the AUGAAU motif overlapping with poly(A) signals"

```
                                                                                          Section 1
                   (1) 1        10        20        30        40        50        60        70            88
             ZM Ubi1 (1) MQIFVKTLTGKTITLEVESSDTIDNVKAKIQDKEGIPPDQQRLIFAGKQLEDGRTLADYNIQKESTLHLVLRLRGGMQIFVKTLTGKT
      S italica Ubi2 (1) MQIFVKTLTGKTITLEVESSDTIDNVKAKIQDKEGIPPDQQRLIFAGKQLEDGRTLADYNIQKESTLHLVLRLRGGMQIFVKTLTGKT
  B distachyon Ubi1 (1) ----------------------------------------------------------------------------------------
 B distachyon Ubi1C (1) ----------------------------------------------------------------------------------------
          Consensus (1) MQIFVKTLTGKTITLEVESSDTIDNVKAKIQDKEGIPPDQQRLIFAGKQLEDGRTLADYNIQKESTLHLVLRLRGGMQIFVKTLTGKT
                                                                                          Section 2
                  (89) 89         100       110       120       130       140       150       160          176
            ZM Ubi1 (89) ITLEVESSDTIDNVKAKIQDKEGIPPDQQRLIFAGKQLEDGRTLADYNIQKESTLHLVLRLRGGMQIFVKTLTGKTITLEVESSDTID
     S italica Ubi2 (89) ITLEVESSDTIDNVKAKIQDKEGIPPDQQRLIFAGKQLEDGRTLADYNIQKESTLHLVLRLRGGMQIFVKTLTGKTITLEVESSDTID
  B distachyon Ubi1 (1) ----------------------------------------------------------------MQIFVKTLTGKTITLEVESSDTID
 B distachyon Ubi1C (1) ----------------------------------------------------------------MQIFVKTLTGKTITLEVESSDTID
         Consensus (89) ITLEVESSDTIDNVKAKIQDKEGIPPDQQRLIFAGKQLEDGRTLADYNIQKESTLHLVLRLRGGMQIFVKTLTGKTITLEVESSDTID
                                                                                          Section 3
                 (177) 177          190       200       210       220       230       240       250         264
           ZM Ubi1 (177) NVKAKIQDKEGIPPDQQRLIFAGKQLEDGRTLADYNIQKESTLHLVLRLRGGMQIFVKTLTGKTITLEVESSDTIDNVKAKIQDKEGI
    S italica Ubi2 (177) NVKAKIQDKEGIPPDQQRLIFAGKQLEDGRTLADYNIQKESTLHLVLRLRGGMQIFVKTLTGKTITLEVESSDTIDNVKAKIQDKEGI
  B distachyon Ubi1 (25) NVKAKIQDKEGIPPDQQRLIFAGKQLEDGRTLADYNIQKESTLHLVLRLRGGMQIFVKTLTGKTITLEVESSDTIDNVKAKIQDKEGI
 B distachyon Ubi1C (25) NVKAKIQDKEGIPPDQQRLIFAGKQLEDGRTLADYNIQKESTLHLVLRLRGGMQIFVKTLTGKTITLEVESSDTIDNVKAKIQDKEGI
        Consensus (177) NVKAKIQDKEGIPPDQQRLIFAGKQLEDGRTLADYNIQKESTLHLVLRLRGGMQIFVKTLTGKTITLEVESSDTIDNVKAKIQDKEGI
                                                                                          Section 4
                 (265) 265 270       280       290       300       310       320       330       340         352
           ZM Ubi1 (265) PPDQQRLIFAGKQLEDGRTLADYNIQKESTLHLVLRLRGGMQIFVKTLTGKTITLEVESSDTIDNVKAKIQDKEGIPPDQQRLIFAGK
    S italica Ubi2 (265) PPDQQRLIFAGKQLEDGRTLADYNIQKESTLHLVLRLRGGMQIFVKTLTGKTITLEVESSDTIDNVKAKIQDKEGIPPDQQRLIFAGK
 B distachyon Ubi1 (113) PPDQQRLIFAGKQLEDGRTLADYNIQKESTLHLVLRLRGGMQIFVKTLTGKTITLEVESSDTIDNVKAKIQDKEGIPPDQQRLIFAGK
B distachyon Ubi1C (113) PPDQQRLIFAGKQLEDGRTLADYNIQKESTLHLVLRLRGGMQIFVKTLTGKTITLEVESSDTIDNVKAKIQDKEGIPPDQQRLIFAGK
        Consensus (265) PPDQQRLIFAGKQLEDGRTLADYNIQKESTLHLVLRLRGGMQIFVKTLTGKTITLEVESSDTIDNVKAKIQDKEGIPPDQQRLIFAGK
                                                                                          Section 5
                 (353) 353  360       370       380       390       400       410       420       430        440
           ZM Ubi1 (353) QLEDGRTLADYNIQKESTLHLVLRLRGGMQIFVKTLTGKTITLEVESSDTIDNVKAKIQDKEGIPPDQQRLIFAGKQLEDGRTLADYN
    S italica Ubi2 (353) QLEDGRTLADYNIQKESTLHLVLRLRGGMQIFVKTLTGKTITLEVESSDTIDNVKAKIQDKEGIPPDQQRLIFAGKQLEDGRTLADYN
 B distachyon Ubi1 (201) QLEDGRTLADYNIQKESTLHLVLRLRGGMQIFVKTLTGKTITLEVESSDTIDNVKAKIQDKEGIPPDQQRLIFAGKQLEDGRTLADYN
B distachyon Ubi1C (201) QLEDGRTLADYNIQKESTLHLVLRLRGGMQIFVKTLTGKTITLEVESSDTIDNVKAKIQDKEGIPPDQQRLIFAGKQLEDGRTLADYN
        Consensus (353) QLEDGRTLADYNIQKESTLHLVLRLRGGMQIFVKTLTGKTITLEVESSDTIDNVKAKIQDKEGIPPDQQRLIFAGKQLEDGRTLADYN

                                                                                          Section 6
                 (441) 441        450       460       470       480       490       500       510           528
           ZM Ubi1 (441) IQKESTLHLVLRLRGGMQIFVKTLTGKTITLEVESSDTIDNVKAKIQDKEGIPPDQQRLIFAGKQLEDGRTLADYNIQKESTLHLVLR
    S italica Ubi2 (441) IQKESTLHLVLRLRGGMQIFVKTLTGKTITLEVESSDTIDNVKAKIQDKEGIPPDQQRLIFAGKQLEDGRTLADYNIQKESTLHLVLR
 B distachyon Ubi1 (289) IQKESTLHLVLRLRGGMQIFVKTLTGKTITLEVESSDTIDNVKAKIQDKEGIPPDQQRLIFAGKQLEDGRTLADYNIQKESTLHLVLR
B distachyon Ubi1C (289) IQKESTLHLVLRLRGGMQIFVKTLTGKTITLEVESSDTIDNVKAKIQDKEGIPPDQQRLIFAGKQLEDGRTLADYNIQKESTLHLVLR
        Consensus (441) IQKESTLHLVLRLRGGMQIFVKTLTGKTITLEVESSDTIDNVKAKIQDKEGIPPDQQRLIFAGKQLEDGRTLADYNIQKESTLHLVLR
                                                                                          Section 7
                 (529) 529  534
           ZM Ubi1 (529) LRGGQ-
    S italica Ubi2 (529) LRGGQ-
 B distachyon Ubi1 (377) LRGGQ-
B distachyon Ubi1C (377) LRGGQ-
        Consensus (529) LRGGQ
```
