## Supplementary-Figure 2 for "Transcription terminator-mediated enhancement in transgene expression in maize: preponderance of the AUGAAU motif overlapping with poly(A) signals"

### 2A

ctgctcgttcagcccacagtaacacgccgtgcgacatgcagatgccctccaccacgccgaccaaccccaagtccgccgcgctcgtccacggcgccatccgcatccgcg
cgtcaacgtcatccggaggaggcgagcgcgatgtcgacggccacggcggcggacacgacggcgacgccccgactccgcgcgcgcgtcaaggctgcagtggcg
tcgtggtggccgtccgcctgcacgagatccccgcgtggacgagcgccgcctccacccagcccctatatcgagaaatcaacggtgggctcgagctcctcagcaacctcc
ccacccccccttccgaccacgctcccttcccccgtgcccctcttctccgtaaacccgagccgccgagaacaacaccaacgaaagggcgaagagaatcgccatagaga
ggagatgggcggaggcggatagtttcagccattcacggagaaatggggaggagagaacacgacatcatacggacgcgaccctctagctggctggctgtcctaaaga
atcgaacggaatcgctgcgccaggagaaaacgaacggtcctgaagcatgtgcgcccggttcttccaaaacacttatctttaagattgaagtagtatatatgactgaaa
tttttacaaggtttttccccataaaacaggtgagcttatctcatccttttgtttaggatgtacgtattatatatgactgaatattttttattttcattgaatgaagattttcgac
cccccaaaaataaaaaacggagggagtacctttgtgccgtgtatatggactagagccatcgggacgtttccggagactgcgtggtgggggcgatggacgcacaacg
accgcattttcggttgccgactcgccgttcgcatctggtaggcacgactcgtcgggttcggctcttgcgtgagccgtgacgtaacagacccgttctcttccccgtctggc
catccataaatccccctccatcggcttccctttcctcaatccagcaccctgattCCGATCGAAAAGTCCCCGCAAGAGCAAGCGACCGATCTCGTG
AATCTCCGTCAAGgtatgcagcctcgcttcctcctcgctaccgtttcaattctggagtaggtcgtagaggataccatgttgatttgacagagggagtagattagat
acttgtagatcgaagtgcggatgttccatggtagatgataccatgttgatttcgattagatcggattaaatctttgtagatcgaagtgcgcatgttccatgaattgcctgt
taccagtagattcaagtttttctgtgttatagaggtgggatctactcgttgagatgattagctcctagaggacaccatgccgttttggaaaatagatcagaaccgtgtag
atcgatgtgagcatgtgttcctgtagatccaagttctttcgcatgttactagttgtgatctattgtttgtgtaatacgctctcgatctatccgtgtagatttcactcgattact
gttactgtggcttgatcgttcatagttgttcgttaggtttgatcgaacagtgtctgaacctaattggatatgtattcttgatctatcaacgtgtaggtttcagtcatgtattta
tgtactccctccgtcccaaattaactgacgtggattttgtataagaatctatacaaatccatgtcagttaattcgggatggagtaccatattcaataatttgtttattgctg
tccacttatgtaccatatgtttgttgttcctcatgtggattctactaattatcattgattggtgatcttctattttgctagtttcctagctcaatctggttattcatgtagatgtg
ttgttgaaatcggagaccatgcttgttattagatagtttattgcttatcagtttcatgttctggttgatgcaacacatattcatgttcgctatctggttgctgcttgatattctc
tgatttacattcattataagaatatattctgctctggttgttgcttctcatgactttacctactcggtaggtgacttaccttttggtttacaattgtcaactatgcag***ATG****ca*
*gatctttgtgaagaccctcaccggcaagaccatcacccttgaggtcgagtcttctgatacgatcgacaatgtcaaggcaaagatccaggacaaggagggcatc*
*cccccggaccagcagcgtctcatcttcgccgggaagcagctggaggatggccgcaccctggcagattacaacatccagaaggagtccaccctccatctggtgct*
*caggctcaggggtggcatgcaaatctttgtgaagacccttactggcaagaccatcacactcgaggtcgagtcgtctgacacgatcgacaatgtgaaggcaaa*
*gatccaggacaaggagggcatcccccccagaccagcagcgcctcatctttgctggcaagcaactggaagacggtcgcaccctggcagattacaatatccagaa*
*ggagtccaccttgcaccttgtgcttcgcctccgtggtggcatgcaaatctttgtcaagaccttgacaggcaagaccattactctggaggtcgagtcgtctgacaca*
*atcgataatgtgaaggcgaagatccaggacaaggagggaattccaccggaccagcagcgcctaatctttgctggcaaacagcttgaggatggccgcaccct*
*ggcagattacaacatccagaaagaatccactctgcacctggtgcttcgcctccgtggtggcatgcagatctttgtcaagaccttgacagggaagacaatcacac*
*tggaggtcgagtcgtctgacacaatcgataatgtgaaggcgaagatccaggacaaggagggcattccaccggaccagcagcgccttatcttcgccggcaagc*
*agcttgaggatggccgcacccttgctgattacaatatccagaaggaatccaccctgcacctggtgcttcgcctccgtggtggcatgcagatctttgtcaagacttt*
*gaccgggaagaccattacactggaggttgaatcttcagacaccatcgacaacgtgaaggcgaagatccaggacaaggagggcatcccccagaccagcag*
*cgcctgatctttgctggtaagcagcttgaggatggacgcactctggcggattataacatccagaaggagtctaccctacacctggtgctccgcctccgtggtggc*
*cag****TAA***gtttgtcaaaaactggcctacagtctgctgcccctgttggtctgccccttggaagtagtcgtgtctatggttatgtgagaagtcgttgtgttctttctaatcccg
tactgtttgtgtgaacatctgctgctgtcgtattgcatcgtgaagaatcctgttatgaataagtgaacatgaaccttgttctgtgattacggcttcgtggttatgcgaacgt
tcttacaaacgcaattgcacctgatgtaaaatcgtttttgctagctgtatggaacaagtgctcatgatgttcatgcaagatgcaattccagcttttgttggtttgtcatcttt
gtactgtgcttaccgcacataaagattgcatcttgcttattgctttgttgctttggtgctcgtccgcttctccttgcaccttatcaaacctttgtttagattctcttcttatagc
acttggtaactctcagctttacaacgccagtactgtttctgaaatttcatgactgataaagctgatagatggagtactaatatatgacatctttccataaatgttcgggtg
cagagatatggaggccccaggatcctatttacaggatgaacctacctgggccgctgtacgcatgacatccgcgagcaagtctgaggttctcaatgtacacatgaaatt
gatttttgctgcgtttggcttggctgatcgttgcatttgttctgattcatcagagttaaataacggatatatcagcaaatatccgcagcatccacaccgaccacacgtccg
gttaacagagtcccccctgccttgctttaattattacggagtactccgctattaatccttagatatgtttcgaaggaactcaaaccttcctccatctgcaaatctcagtgctt
caaaactggaattagataattgaaaccttcattcggttgcaattcacaactgcaaattgaacagcactgtcaatttcaatttcgggttcacgattccaccgataggttga
catgatccatgatccacccattgtacaac

**2B**

ggcgtcaggactggcgaagtctggactctgcagggccgaactgctgaagacgaagcagaggaagagaaagggaagtgttcgacttgtaatttgtaggggtttttttta
gaggaacttgtaatttgtaggtgggctggcctcgttggaaaaacgatgctggctggttgggctgggccgatgtacgcttgcaaacaacttgtggcggcccgttctggac
gagcaggagtttctttttgttctcacttttctggtcttctttagttacggagtaccttttgtttttaaaggagttaccttttttttaggaattctttagttacctttcgcttgctc
tcaaaaaatatttaactttcgctttttttcattttaatttttgcaactatttacgagtttcatgaatgcttattttccagcatatcattatttgcaagtatttttatgccgtatgt
attggacgagagccatcgggactgttccagagactgcgtggtggggacggctcccaaccgccttttctatctctgttcgcatccggtggccgacttggctcgcgcgtga
gccgtgacgtaacagacttggtctcttccccatctggccatctataaattcccccatcgatcgaccctccctttccCCAATCCAGCACCCCCGATCCCGATCG
AAAATTCTCCGCAACAGCAAGCGATCGATCTAGCGAATCCCCGTCAAGgtatgtagcctctcgattcctcctcagccctgccctcgatttggtgta
cgcgttgagatgatgatctcgtagatgtctagatgacaccatgtcgatttgaaatagatcagatccgtgtagatcgatgagctcctgtgtacctgtggattcaagttattt
tcgcatgctattgttgtgatctactagatctagtgtgtgtattctatgctatcgatttctccgtgtagatttcactcgattactgttactgtggcttgatcggccatagatgtt
ggttaaggtttgatcggttagtgtttgaacctgcgtggatatctagcatccatctattatcgtgtaggtttcgaacaaacaagcactattattgtactgatggttcgtctat
ggttggttttgaccgttttagtgttgaacgagccttctgtatttgtttattgctgtccagtgatgtaccatgttcgttgagtgtcggattatactaattattgttgattgataat
cttgtagtttgcttttcctaatttatttatcgtagtcctgatttgcctcagctgtgcctcacccgtgcgatggtcaatcaacttgttagcccaatctgcttaatcatgtacattt
gttgttagaatcagagatcaagccaattagctatcttattgcttatctgttccatgttctgatcgatgtaacagtctacactttgctctgtgctacttgattaaaacattct
gacttaaattcatgattggaagtttcagatctgattgttgccttacttgactaatatctattcatgtgacacctctctgtcttggtaacttaccgctgtttgtttgtaatttctg
actatgcag***ATG****cagatctttgtgaagaccctcactggcaaaaccatcacccttgaggtcgagtcgtccgacacgatcgacaacgtcaaggcaaagatccag*
*gacaaggagggcattcctccagaccagcagcgcctcatctttgctggaaagcagcttgaggacggccgcaccctcgccgactacaacatccagaaggagtcc*
*accctccacctggtcctgaggctccgtggtggcatgcagatcttcgtcaagacccttaccggcaagaccatcacgctggaggtcgagtcctctgacacgatcgac*
*aatgtgaaggcgaagatccaggataaggagggcatcccccggaccagcagcgcctcatctttgccggcaagcagcttgaggacggccgtaccctcgccgac*
*tacaacatccagaaggaatccacactccatctggtgctcaggctgcgtggtggcatgcagatcttcgtcaagaccctaaccggcaagaccatcactctggaggt*
*tgagtcctctgacacgatcgacaatgtgaaggcaaagatccaggataaggagggcattcccccggaccagcagcgcctcatcttcgctggcaagcagcttga*
*ggatggccgcaccctggcagattacaatatccagaaggaatccaccttgcacctggtgcttcgcctccgtggtggcatgcagatctttgtaaagaccttgactgg*
*caagacaattaccctggaggttgagtcgtccgacacaattgacaatgtcaaggcgaagatccaggacaaggagggcatcccaccggaccagcagcgcctca*
*tcttcgccggcaagcagcttgaggatggtcgcacccttgcagattacaatatccagaaggaatccactctgcatctggtgcttcgtctccgcggtggaatgcaga*
*tcttcgttaagacgttgacagggaagaccatcacactggaggttgaatcttcggacaccattgacaacgtgaaggcaaagatccaggacaaggagggcatcc*
*ccccagaccagcagcgcctcatctttgctggtaagcagcttgaggatggccgcacccttgcagattacaacattcagaaggagtccaccctgcacctggtgctcc*
*gtctccgtggtgggcag****TAA***gcttctgccgaactggttcacagtctgctgcccttggtggtctgccccttagtggtcatgccttttgttatgtgtcttgcgtcccaatcct
gtatcgtttgtgtgaacatctctgctgctgtatagcagcttgaatcctgttatgaatttgtgaacctgaaccttgttccgtgaatcatgttatgaataagtgaacctgaacc
ttgttccgtgattattgttacaatctgttgtgccgtatggttggtcgtgtgtgatttatgttgaactggagaaccaagttcgttccaggacatattgcaacctaagctaaac
catgtagaactacttgttctgggagacataaaacgtcatttttatgcattcgtaacatttaagcatactacaataattgtattgtccttttcctactcatccttgaaaccat
atgcctcttctcagcgcctctacatgcagtgtgctcagaacaaacaggccctgccagctgcttttcaattttccaattaataaccacaatagtcggactatggcatctgtg
ggtgactatgcaagatgttgctgtcaggtctctgaaacttttcccatgtatctgttgaaattacccagtaaattcatgcctctatttaatctggcatggttgattttcaaac
agaatgtgttttttttgttctggaagctatattggtaaataaatacaaagctggagtgtgattatatttccaacagatattcaagaaaatctcagttgatttatttactac
tgtagtatatatatatatcttacagttgacttctcatatttcaaacgacatgtgagcacattgttcagtttcttaggatgtgttgtgtgctcaaaggtgtaattttgcattctg
ccctccgagtaaacactacacgtatttttttgagtggcagtgcatttgattacaaggcaacaacaacaaaaacctatggcaagatatccttcttagaggctgccaggat
cattttgactgaactatgtaaggctgaagaaaagg

**2C**

tgcgtctggacgcacaagtcatagcattatcggctaaaatttcttaatttctaaattagtcatatcggctaagaaagtggggagcactatcatttcgtagaacaagaac
aaggtatcatatatatatatatatataatatttaaactttgttaagtggaatcaaagtgctagtattaatggagtttcatgtgcattaaattttatgtcacatcagcaattt
tgttgacttggcaaggtcatttagggtgtgtttggaagacaggggctattaggagtattaaacatagtctaattacaaaactaattgcacaaccgctaagctgaatcgc

gagatggatctattaagcttaattagtccatgatttgacaatgtggtgctacaataaccatttgctaatgatggattacttaggtttaatagattcgtctcgtgatttagcc
tatgggttctgctattaattttgtaattagctcatatttagttcttataattagtatccgaacatccaatgtgacatgctaaagtttaaccctggtatccaaatgaagtctta
tgagagtttcatcactccggtggtatatgtacttaggctccgttttcttccaccgacttatttttagcacccgtcacattgaatgtttagatactaattagaagtattaaacg
tagactatttacaaaatccattacataagacgaatctaaacggcgagacgaatctattaaacctaattagtccatgatttgacaatgtgttgctacagtaaacatttgct
aatgatggattaattaggcttaatagattcgtctcgccgtttagcctccacttatgtaatgggttttctaaacaatctacgtttaatactcctaattagtatctaaatattca
atgtgacacgtgctaaaaataagtcagtggaaggaagagaacgtccccttagttttccatcttattaattgtacgatgaaactgtgcagccagatgattgacaatcgca
atacttcaactagtgggccatgcacatcagcgacgtgtaacgtcgtgagttgctgttcccgtagAGAAATATCAACTGGTGGGCCACGCACATCAGCG
TCGTGTAACGTGGACGGAGGAGCCCCGTGACGGCGTCGACATCGAACGGCCACCAACCACGGAACCACCCGTCCCCACCTCTC
GGAAGCTCCGCTCCACGGCGTCGACATCTAACGGCTACCAGCAGGCGTACGGGTTGGAGTGGACTCCTTGCCTCTTTGCGCTG
GCGGCTTCCGGAAATTGCGTGGCGGAGACGAGGCGGGCTCGTCTCACACGGCACGGAAGACgtcacgggttccttccccacctctcctc
ttccccaccgccataaatagCCGACCCCCTCGCCTTTCTCCCCAATCTCATCTCGTCTCGTGTTGTTCGGAGCACACCACCCGCCCCAAA
TCGTTCTTCCCGCAAGCCTCGGCGATCCTTCACCCGCTTCAAGgtacggcgatcgtcttcctcctctagatcggcgtgatctgcaagtagttgatttg
gtagatggttaggatctgtgcactgaagaaatcatgttagatccgcgatgtttctgttcgtagatggctgggaggtggaatttttgtgtagatctgatatgttctcctgttt
atcttgtcacgctcctgcgatttgtggggattttaggtcgttgatctgggaatcgtggggttgcttctaggctgttcgtagatgaggtcgttctcacggtttactggatcatt
gcctagtagatcagctcgggctttcgtctttgtatatggtgcccatacttgcatctatgatctggttccgtggtgttacctaggtttctgcgcctgattcgtccgatcgatttt
gttagcatgtggtaaacgtttggtcatggtctgatttagattagagtcgaataggatgatctcgatctagctcttgggattaatatgcatgtgtcaccaatctgttccgtg
gttaagatgatgaatctatgcttagttaatgggtgtagatatatatgctgctgttcctcaatgatgccgtagcttttacctgagcagcatggatcctcctgttacttaggta
gatgcacatgcttatagatcaagatatgtactgctactgttggaattctttagtatacctgatgatcatccatgctcttgttacttgttttggtatacttggatgatggcatg
ctgctgctttttgttgatttgagcccatccatatctgcatatgtcacatgattaagatgattacgctgtttctgtatgatgccatagcttttatgtgagcaacatgcatcctc
ctggttatatgcattaatagatggaagatatctattgctacaatttgatgattattttgtacatacgatgatcaagcatgctcttcatactttgttgatatacttggataatg
aaatgctgctgcacgttcattctatagcactaatgatgtgatgaacacgcacgacctgtttgtggcatctgtttgaatgtgttgttgctgttcactagagactgttttatta
acctactgctagatacttacccttctgtctgtttattcttttgcag***ATG****cagatctttgtcaagaccctcaccggcaagaccatcaccctcgaggtggagtcttctgac*
*accattgacaacgtcaaggccaagatccaggacaaggaaggcattcccccggatcagcagcggctcatctttgccggcaagcagcttgaggatgggcgcacc*
*ctggctgactacaacatccagaaggagagcaccctccacctggtgctccgtctcaggggaggcatgcagatctttgtgaagaccttgactggcaagaccatca*
*cccttgaggtggagtcttccgacaccatcgacaacgtcaaggccaagatccaggacaaggagggcatcccccggaccagcagaggctcatctttgcgggca*
*agcagcttgaggatggacgcaccctggctgactataacatccagaaggagagcaccctccatctggtgctccgtctcaggggaggcatgcagatcttcgtgaa*
*gactctcactggcaagaccatcaccctcgaggtggagtcttccgacaccatcgacaacgtcaaggccaagatccaggacaaggagggcatcccccccagacca*
*gcagaggctcatctttgctggcaagcagcttgaggacggacgcaccctggctgactataacatccagaaggagagcaccctccacctggtgctccgcctgagg*
*ggtgggatgcagatctttgtgaagactttgactggcaagaccattactttggaggttgagagctccgacaccatcgacaacgtgaaggccaagatccaggac*
*aaggaaggcatccccccggaccagcagaggctcatcttcgccggcaagcagcttgaggacggacgcaccctggctgactataacatccagaaggagagcac*
*cctccacctggtgctccgtctcaggggaggcatgcagatcttcgtgaagaccctcactggcaagaccatcacccttgaggtggagtcttccgacaccatcgacaa*
*tgtcaaggccaagatccaggacaaggagggcatcccccccagaccagcagagactcatctttgcaggcaagcagcttgaggacggacgcaccctggctgact*
*acaacatccagaaggagagcaccctccacctggtgctccgtctcaggggaggcatgcagatcttcgtgaagaccctcactggcaagaccatcaccctcgaggt*
*ggagtcttctgacaccatcgacaacgtcaaggccaagatccaggacaaggagggcatcccccggaccagcagcgtcttatctttgccggcaagcagctgga*
*ggatggccgcacccttgcggattacaatatccagaaggagagcaccctccatctggtgctccgtctgaggggtgggatgcagatattcgtgaagactttgaccg*
*gcaagaccatcactttggaggttgagagctccgacaccattgacaatgtgaaggccaagatccaggacaaggagggtatccccccggaccagcagcgtctg*
*atcttcgccggcaagcaactggaggatggccgcaccctggcggactacaatatccagaaggagtccaccctccacctggtgctccgcctccgtggtggtcag****T***
***AA***gcccatcggtcatggatgcttctactgtacctgggtcgtctggtctctgcctgtgtcacctttgaagtacctgtgtcgggattgtgtttggtcatgaactgcagtttgtc
tttgatgttcttttgtctggtcttatgaactggttgtatctgtatgtttactgtaaactgttgttgcggtgcagcagtatggcatccgaatgaataaatgatgtttggactta
aatctgtactctgtttgttttcggttatgccagttctatattgcctgagatcagaatgtttagcttttgagttctgtttggcttgtggtcgactcctgtttcttacttgaggcgt
aactctgttctggcaaactcaaatgtctaactgaatgttttaggacttaattgttggacagattaacgtgtttggtttgtttctagattgtgattcggaaggcttgttagttg
tggaatcaaggagagcagctaggtctgtgcagaacgttattttggatttaagcctttctcagattatgccattactctaaacctaatgatatcatatttcactcggggatgt
tggagtagtcttttctttctcctgcagacaaaatgattttgctttcgtgtgtgtacatgattttgtgcaactgttgcaacaactgaagtagacaagttttgacctcaccaga
agaatgaaaaagattttggaatttgttacatcgacaaaccattgtaacttggcccatcagaatgcacagaagagcggctacaaattgacatgcgttgcaaactttgca
atagttgatgcacatgtttgccattgcctgccagtcttaggaaaagtgtgtggttcgagaaatctaagcatatgtgctctgctcacattgcgtggaacccacacagcttt

gtcacactcttgtccactccagaagtcattcctggcgctgtttacccctggtaaaaggtaaccgaaaacttctcaaggctgtacccaaaactggaaggaaatttggagg
aaatctttgcttttgatcggctcactctttc
